# The attentive reconstruction of objects facilitates robust object recognition

**DOI:** 10.1101/2023.07.17.549340

**Authors:** Seoyoung Ahn, Hossein Adeli, Gregory J. Zelinsky

## Abstract

Humans are extremely robust in our ability to perceive and recognize objects—we see faces in tea stains and can recognize friends on dark streets. Yet, neurocomputational models of primate object recognition have focused on the initial feed-forward pass of processing through the ventral stream and less on the top-down feedback that likely underlies robust object perception and recognition. Aligned with the generative approach, we propose that the visual system actively facilitates recognition by reconstructing the object hypothesized to be in the image. Top-down attention then uses this reconstruction as a template to bias feedforward processing to align with the most plausible object hypothesis. Building on auto-encoder neural networks, our model makes detailed hypotheses about the appearance and location of the candidate objects in the image by reconstructing a complete object representation from potentially incomplete visual input due to noise and occlusion. The model then leverages the best object reconstruction, measured by reconstruction error, to direct the bottom-up processing of selectively routing low-level features, a top-down biasing that captures a core function of attention. We evaluated our model using the MNIST-C (handwritten digits under corruptions) and ImageNet-C (real-world objects under corruptions) datasets. Not only did our model achieve superior performance on these challenging tasks designed to approximate real-world noise and occlusion viewing conditions, but also better accounted for human behavioral reaction times and error patterns than a standard feedforward Convolutional Neural Network. Our model suggests that a complete understanding of object perception and recognition requires integrating top-down and attention feedback, which we propose is an object reconstruction.

**Author Summary:** Humans can dream and imagine things, and this means that the human brain can generate perceptions of things that are not there. We propose that humans evolved this generation capability, not solely to have more vivid dreams, but to help us better understand the world, especially when what we see is unclear or missing some details (due to occlusion, changing perspective, etc.). Through a combination of computational modeling and behavioral experiments, we demonstrate how the process of generating objects—actively reconstructing the most plausible object representation from noisy visual input—guides attention towards specific features or locations within an image (known as functions of top-down attention), thereby enhancing the system’s robustness to various types of noise and corruption. We found that this generative attention mechanism could explain, not only the time that it took people to recognize challenging objects, but also the types of recognition errors made by people (seeing an object as one thing when it was really another). These findings contribute to a deeper understanding of the computational mechanisms of attention in the brain and their potential connection to the generative processes that facilitate robust object recognition.

## 1 Introduction

One of the hallmarks of human visual recognition is its robustness against various types of noise and corruption (Avidan et al., 2002; Vogels & Biederman, 2002). Convolutional Neural Networks (CNNs) trained on object classification have gained popularity as models of human and primate vision due to their ability to predict behavioral and neural activity during visual recognition (Cadieu et al., 2014; Cichy, Khosla, Pantazis, Torralba, & Oliva, 2016; Geirhos et al., 2021; Kriegeskorte, 2015; Schrimpf et al., 2020). However, CNNs can still be highly vulnerable to small changes in object appearances even when the changes are almost imperceptible to humans (Dodge & Karam, 2017; Szegedy et al., 2013; but see Geirhos et al., 2021 for CNNs reaching the human-level performance in recognizing distorted images). Recent studies also revealed that CNNs exploit fundamentally different feature representations than those used by humans (e.g., relying on texture rather than shape, Baker, Lu, Erlikhman, & Kellman, 2018; Geirhos, Rubisch, et al., 2018), suggesting that current computational vision models still do not fully capture primate visual processing.

The question of how humans robustly recognize objects under challenging viewing conditions, such as noise and occlusion, has been a long-standing problem spanning various disciplines, from cognitive science (Biederman, 1987; Tarr & Pinker, 1990) and neuroscience (Plaut & Farah, 1990; Rolls, 1994) to computer vision (Marr, 1982; Ullman, 1989). One widely accepted hypothesis is that top-down feedback, broadly associated with attention control or bias, serves to refine object representations through the selective modulation of the responses in early visual areas, thereby better aligning them with higher-level object hypotheses (Bar et al., 2006; Carpenter & Grossberg, 1987; Gilbert & Li, 2013; Lee & Mumford, 2003; Ullman, 1995). However, the computational mechanism underlying this top-down feedback remains elusive, particularly regarding how this modulation can selectively target specific locations and features of visual stimuli, as demonstrated in the experimental literature (Martinez-Trujillo & Treue, 2004; Müller & Hübner, 2002; Posner, 1980; Treisman, 1988). This challenge is particularly complex considering that object representations in higher cortical areas are largely invariant to changes in the lower-level representation of an object’s location or features (DiCarlo, Zoccolan, & Rust, 2012). In a recent perspective on the modeling of object-based attention (Cavanagh et al., 2023), a failure to explain how coarse object representations from high-level cortex effectively modulate lower-level feature or spatial attention was identified as a critical shortcoming of current approaches.

Our approach is to apply generative modeling to this question. A generative model of perception suggests that the brain constantly attempts to generate percepts consistent with our hypotheses about the world (Dayan, Hinton, Neal, & Zemel, 1995; de Lange, Heilbron, & Kok, 2018; Yuille & Kersten, 2006), a process referred to as “prediction” in predictive coding theory (Clark, 2013) or “synthesis” in analysis-by-synthesis theory (Yuille & Kersten, 2006). Although direct experimental evidence for these processes in the brain remains limited, various visual phenomena strongly suggest the involvement of top-down generative or reconstructive processes in shaping human visual perceptions (e.g., visual imagery, object completion, and pareidolia). We propose a new model of object recognition that generates an object reconstruction—a visualized prediction about the possible appearance of an object—and uses it as top-down attention feedback to bias feed-forward processing. This model, called Object Reconstruction-guided Attention (ORA), is based on the fundamental premise that the process of object reconstruction serves as the underlying mechanism through which high-level object representations are used to recover specific location and feature information. This enables the model to exert precise top-down biasing control over both spatial and feature dimensions, a function typically associated with separate attention systems (Carrasco, 2011; Cavanagh et al., 2023; Kravitz & Behrmann, 2011). Under ORA, top-down biasing occurs by prioritizing the object features that offer the most accurate interpretation of the retinal input, that is, minimizing the reconstruction error (similar to “explain-away”, Clark, 2013). Although promising recent work has modeled human visual cortical processing using a neural network equipped with a decoder to reconstruct the input (Adeli, Ahn, & Zelinsky, 2023; Al-Tahan & Mohsenzadeh, 2021; Boutin, Zerroug, Jung, & Serre, 2020; Csikor, Mesźena, Szabó, & Orbán, 2022; Fleming & Storrs, 2019; Hedayati, O’Donnell, & Wyble, 2022; Xing, Chrastil, Nitz, & Krichmar, 2022; Yildirim, Belledonne, Freiwald, & Tenenbaum, 2020), no current models use top-down reconstruction feedback to improve the robustness of object perception and recognition.

ORA fills this gap by providing a new computational model that integrates object recognition and attention by leveraging object reconstructions to generate top-down attention feedback. As part of its novel neural network architecture, the model directs more attention to the areas and features of the input that align with the current best object reconstruction. This targeted attention effectively enhances otherwise weak bottom-up input signals, thereby reducing the interference from noise and occlusion and improving recognition performance. In extensive model evaluation, we found that ORA exhibits remarkable robustness as evidenced by its effective mitigation of noise and occlusion effects when applied to challenging recognition tasks such as MNIST-C (handwritten digits under corruptions, Mu & Gilmer, 2019) and ImageNet-C (real-world objects under corruptions, Hendrycks & Dietterich, 2019). ORA also accounts for behavioral reaction time and error results that we observed from human participants performing the same tasks, better than a baseline feed-forward network (e.g., a ResNet; He, Zhang, Ren, & Sun, 2016). We hope that ORA’s use of object-centric representation and reconstruction will provide new insights into the neural mechanism endowing humans with robust object perception and recognition.

## 2 Object Reconstruction-guided Attention

The concept of interactive top-down and bottom-up processing is a central theme in numerous influential cognitive neuroscience and vision theories (Bar et al., 2006; Carpenter & Grossberg, 1987; Dayan et al., 1995; Friston, 2010; Gilbert & Li, 2013; Lee & Mumford, 2003; Ullman, 1995; Yuille & Kersten, 2006). Among these theories, our model builds most closely upon Grossberg and Carpenter’s Adaptive Resonance Theory (ART). ART constructs expected category representations or *templates*, which it uses via top-down feedback to inhibit bottom-up features that do not align with these expectations (referred to as “top-down attentive matching”, Grossberg, 2017). Our model, ORA, advances the fixed categorical template approach of ART by incorporating a generative mechanism that dynamically selects the most probable object template based on the input, achieved through object reconstruction. This approach enables ORA to adaptively adjust its top-down expectations to accommodate variability in exemplars and noise in the visual input, thereby making it robust. ORA also demonstrates that top-down attentive biases discussed in the cognitive science literature, such as object-fitting attentional shrouds (Fazl, Grossberg, & Mingolla, 2009; Müller & Hübner, 2002) and the formation of robust object entities via attentive feature binding (Hummel & Biederman, 1992; Treisman, 1988), do not require explicit learning. Instead, these biases may emerge via the active reconstruction and use of hypothesized objects to guide bottom-up object recognition, as theorized by ORA.

Figure 1 shows how ORA might be implemented in the brain. The retinal image is first processed through the ventral stream, which consists of multiple layers of linear and non-linear filters (e.g., V1, V2, and V4) specialized in extracting the visual features of objects (DiCarlo et al., 2012). As information flows through these layers, neuronal receptive field sizes progressively increase and responses grow more selective and abstract (depicted by the greenish circles). For our implementation, we model the ventral stream using a Convolutional Neural Network (CNN) incorporating skip connections between retinotopic feature maps (He et al., 2016). At the top of this hierarchy is the inferotemporal cortex (IT), where the low-dimensional latent features are encapsulated into separate category-selective slots (Rajalingham & DiCarlo, 2019)^1^. Unlike conventional CNNs and neural networks, which typically assign a single output neuron to represent each category, these object slots employ a vector representation in which a set of neurons represents each category and each neuron within this set encodes meaningful variations along specific feature dimensions of objects (e.g., scale, thickness, etc). This allows the model to construct robust yet flexible object representations (Peters & Kriegeskorte, 2021; Sabour et al., 2017) that maintain selectivity for the distinctive features of objects while being tolerant to variability in these features, consistent with object file theory (Kahneman et al., 1992).

**Fig 1.**
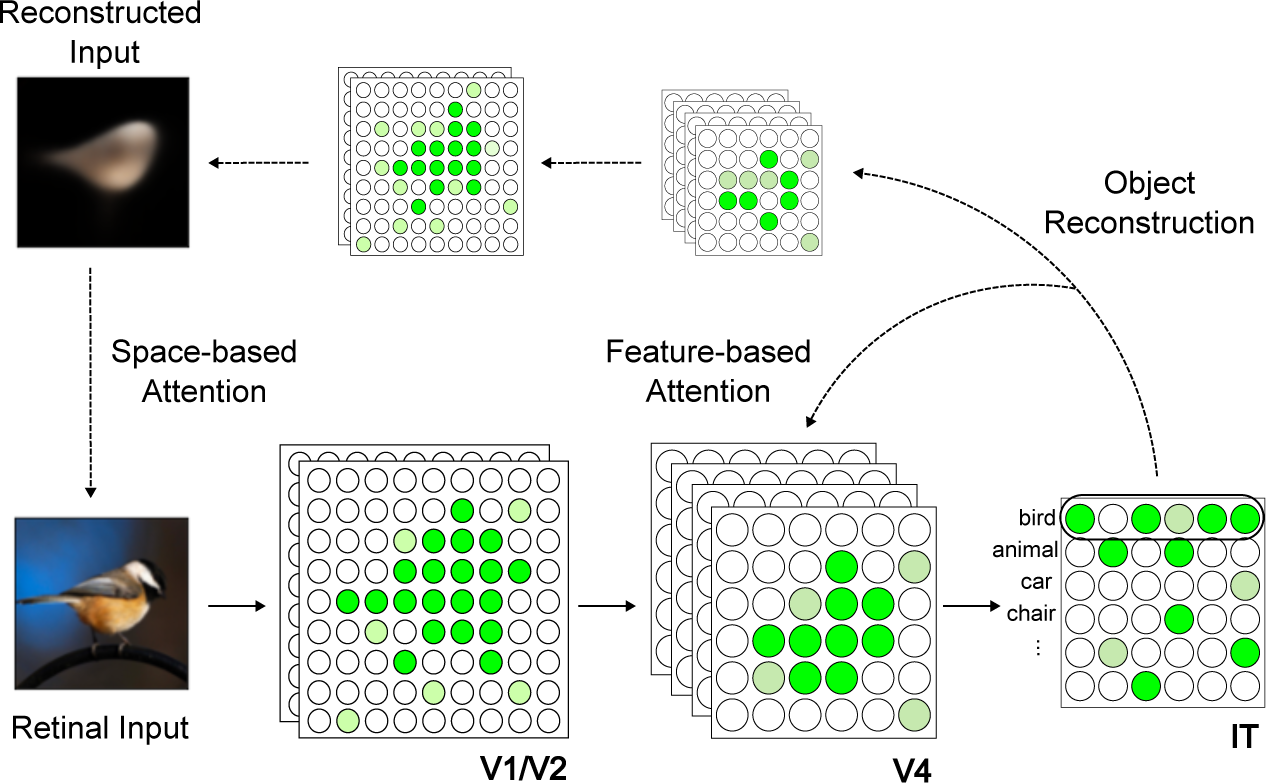
A potential implementation of ORA in the brain. ORA uses object reconstruction as top-down attention feedback and iteratively routes the most relevant visual information in successive steps of a feed-forward object recognition process. This reconstruction-guided attention operates at two levels, 1) Spatial attention: a long-range projection that spatially constrains attention to the most likely object location and 2) Feature-based attention: more local feedback that biases feature binding strengths to favor the formation of a better object reconstruction. See main text for more details.

The object features that ORA encodes into its IT slots are then decoded to generate top-down reconstructions of objects in the visual input. This encoding-decoding process is similar to an autoencoder architecture (Hinton & Salakhutdinov, 2006), where a model first encodes a retinal input to a highly compressed representation and then expands this latent representation through a decoder to reconstruct the original input. ORA’s decoding process is different in that it iteratively generates reconstructions (rather than generating only one) and uses the early reconstructions to recover a representation of retinotopy and to facilitate the binding of object features to specific locations. ORA decodes an object reconstruction using dedicated feedback connections from an inversely mirrored CNN (Dumoulin & Visin, 2018). This CNN decoder takes the most activated object slot in ORA’s IT cortex (measured by the sum of squares of all neural activation within that slot, i.e., vector magnitude), and sequentially reconstructs the input features using a series of convolutional and unpooling layers. More details about ORA’s architecture can be found in the Methods.

ORA relies on a neural network trained with two objectives: accurately classifying an input and generating its reconstruction. Margin loss (Sabour et al., 2017) was used for classification training. By minimizing this loss, the activation of the target object slot (measured by vector magnitude) is pushed toward the desired value of 1, while the activation of non-target object slots is suppressed to near 0 (see Methods). We compute reconstruction loss as the mean squared differences in pixel intensities between the input image and the model’s reconstruction. During model evaluation (i.e., inference), ORA leverages object reconstructions as top-down attentional templates, selectively routing the bottom-up processing of features that align with top-down expectations, similar to ART’s iterative “matching” process (Carpenter & Grossberg, 1987; Grossberg, 2017). By reconstructing the details of object appearances, ORA can exert nuanced attention control over various levels of visual features, both spatial and non-spatial, indicated by the downward arrows in Figure 1. The iterative process of top-down matching and bottom-up routing in ORA continues until the network stabilizes, determined by the model’s classification confidence surpassing a predefined threshold. See Section 5.5 of the Methods for details about how this confidence was estimated.

### 2.1 Reconstruction-based spatial masking

Our goal in this study is to show that ORA’s reconstruction-guided feedback can serve two significant roles with respect to attention. The first of these is the control of spatial attention. Spatial attention refers to the prioritization of information at a spatial location by filtering or masking out information from all other locations (Posner, 1980), with the neural expression of this being enhanced activity at the attended location and inhibition elsewhere. Several experimental studies have reported that spatial attention can dynamically configure itself to align with an object’s shape (Blaser, Pylyshyn, & Holcombe, 2000; Chen & Zelinsky, 2019; Moore & Fulton, 2005; Müller & Hübner, 2002), and this has even been modeled as object-fitting attentional shrouds (Fazl et al., 2009; Tyler & Kontsevich, 1995). ORA posits that the visual system constructs an attentional shroud around an object by iteratively comparing a model generated object reconstruction with incoming sensory signals. This iterative process involves generating a spatial priority mask, a function known to be mediated by the dorsal pathway encompassing posterior parietal cortex (PPC; Behrmann, Geng, & Shomstein, 2004; Xu, 2018), which we implement through simple binary thresholding. The resulting spatial mask is then utilized to modulate activity in early ventral areas, suppressing neural responses to features outside the expected object shape. ORA repeated this process until the model’s classification confidence surpassed a predetermined threshold or until the limit of five forward processing steps was reached, whichever came first (see Methods for additional details). Figure 2 demonstrates how the reconstruction-based spatial masking effectively filters out background noise, thereby enhancing classification accuracy. Initially, ORA hypothesized that the input was the digit 5, but after filtering out noise using its reconstruction mask the model correctly changed its hypothesis to a 3, matching the ground truth. On average, ORA’s iterative spatial masking enabled it to successfully recover the object’s correct identity on over 90% of the trials when the model started with an incorrect hypothesis (see Extended Data S1). Previous research has similarly argued for benefits from a reconstruction-based masking approach, but in the context of augmenting datasets for improving the learning of robust visual representations for use in downstream tasks (He et al., 2022; Sabour et al., 2017). ORA is different from previous modeling work (Adeli et al., 2023; Deco & Rolls, 2004; Jeurissen, Self, & Roelfsema, 2016) in that it leverages object reconstructions learned during training to generate spatial attention masks during inference. These masks therefore bias model processing to the hypothesized shape of the object.

**Fig 2.**
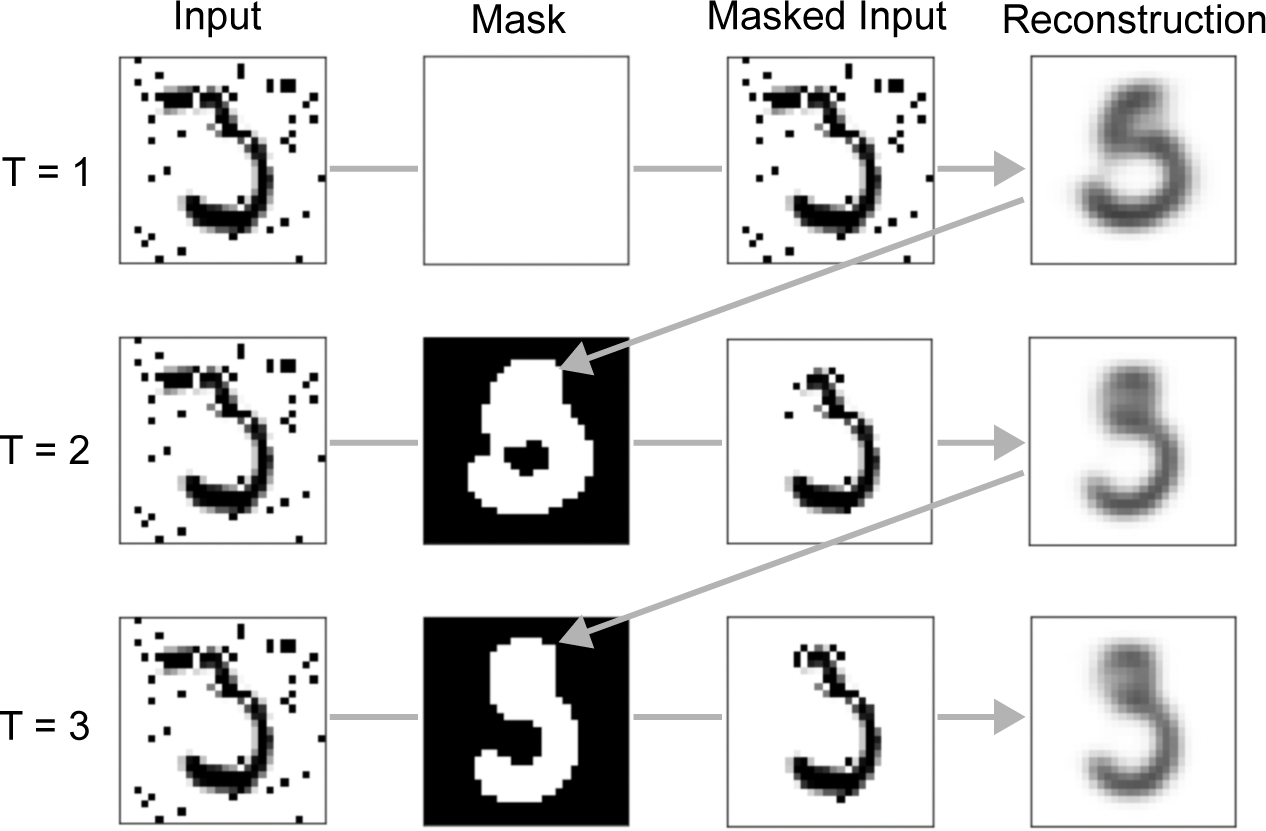
Object reconstructions and generated spatial masks for three forward processing steps (out of a possible five in total). The spatial mask serves to limit ORA’s attention to the shape of the most likely object. In this example, the model’s most likely digit hypothesis changed from a 5 to a 3. The groundtruth class is 3.

### 2.2 Reconstruction-based feature binding

A second role of attention served by ORA’s reconstruction-based top-down biasing is the binding of features into objects (Greff et al., 2020). Feature binding refers to the process by which the varied features of an object are spatially integrated to form a coherent object representation (e.g., horizontal, vertical, and curved lines are bound top-to-bottom to form the digit 5), and this process is believed to be fundamental to robust object perception and recognition (Fazl et al., 2009; Hummel & Biederman, 1992; Treisman, 1988). Recent advances in computer vision demonstrated that training models to reconstruct objects resulted in robust feature binding enabling the grouping and tracking objects within a scene (Greff et al., 2020; Locatello et al., 2020; Sabour et al., 2017). ORA accomplishes feature binding by leveraging an object reconstruction to determine the features that should be associated with specific objects. Specifically, ORA modulates the weights between the V4 and IT layers, suppressing the binding strength between a feature and an object if it leads to a poorer reconstruction match to the input (see Methods for pseudocode). By selectively suppressing information incongruent with ORA’s top-down object reconstruction, this algorithm progressively refines neural selectivity to converge towards the most probable object hypothesis, thereby narrowing the population response. ORA’s binding mechanism is consistent with the information tuning model of feature-based attention (Carrasco, 2011; Martinez-Trujillo & Treue, 2004), and it can be interpreted as a form of biased competition where the bottom-up control for neural resources (i.e., receptive fields) is biased by a higher-level representation of an object (Beck & Kastner, 2009; Bundesen, Habekost, & Kyllingsbæk, 2011; Desimone & Duncan, 1995). Figure 3 shows ORA’s binding coefficients (which modulate the original weights between the V4 and IT layers) sharpening over iterations and becoming more focused on the specific object features that best match the input. Binding coefficients were updated over three iterations during inference (testing); coefficients were not updated during model training. Available computing resources required us to limit ORA’s reconstruction-based feedback to only three iterations, however we observed during piloting that three steps was usually sufficient for ORA to converge on a classification.

**Fig 3.**
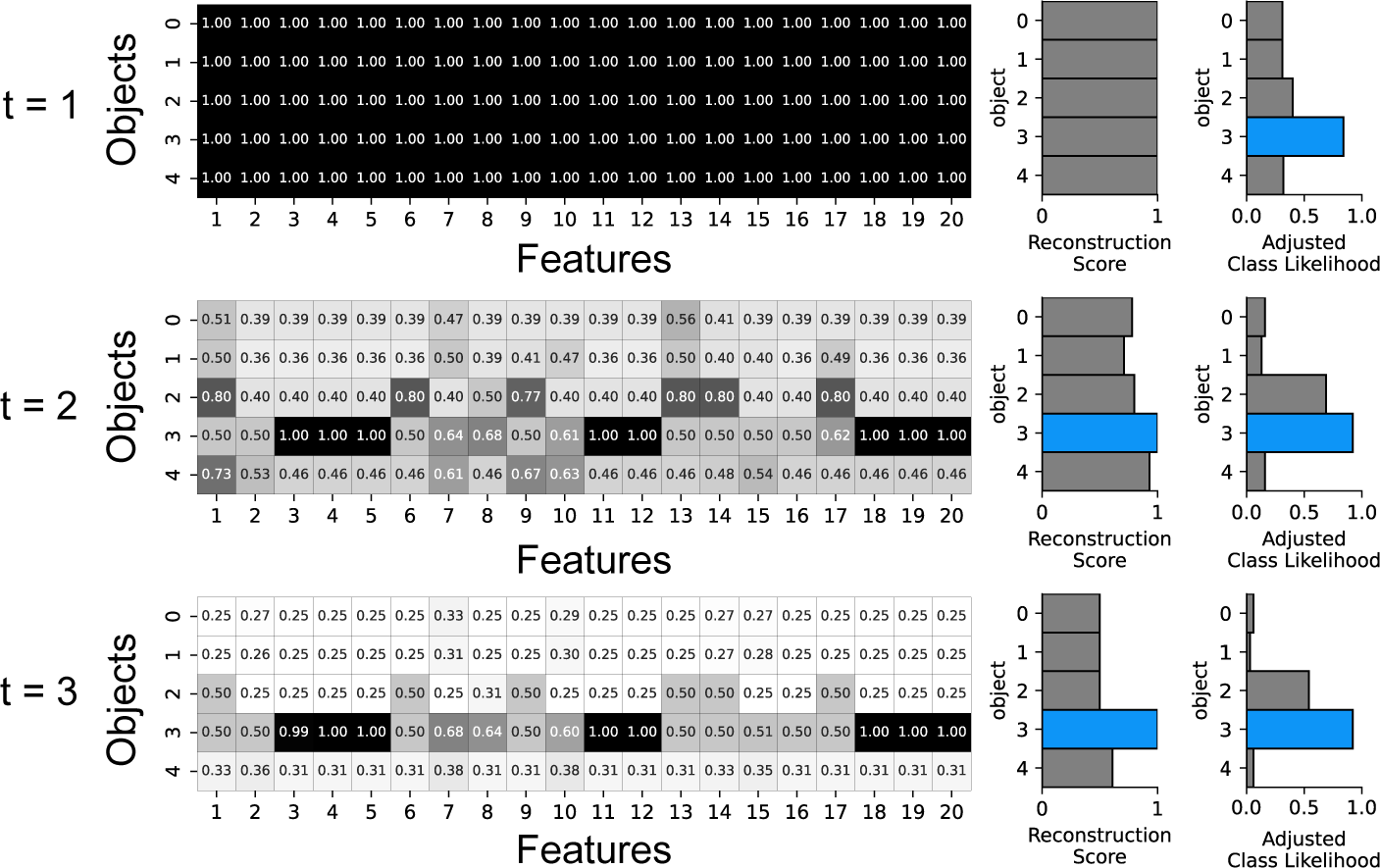
Step-wise visualizations of reconstruction-based feature binding. The matrices show binding coefficients between object slots (rows) and feature slots (columns) for three time steps. The binding coefficients were initialized to one at t = 1, resulting in the dark matrix on the top, but in the subsequent time steps there is a rapid selective suppression of coefficients (matrix cells becoming lighter) as the model learns the object features leading to higher reconstruction accuracy. In the illustrated example, the coefficient matrix becomes sparser with each iteration as ORA focuses its attention on the features of the digit 3 (darker line forming along the fourth row). The middle column of bar graphs show the hypothesis yielding the highest reconstruction score (in blue), and the rightmost column shows the class likelihood adjusted by the reconstruction score. For clarity, only 5 object slots and 20 feature slots are illustrated, but the full figure can be found in Extended Data S2.

## 3 Results

We evaluated ORA in challenging object recognition tasks. First, we compared ORA’s recognition performance to that of a standard feed-forward CNN using a version of the MNIST dataset designed to test for effects of occlusions and specific forms of visual noise (MNIST-C; Mu & Gilmer, 2019). Both ORA and the CNN were trained on “clean” handwritten digits and then systematically tested on each form of corrupted digit. This evaluation revealed that ORA used its reconstruction-based feedback to filter out irrelevant information and this led to greater recognition robustness to the local changes in texture introduced by the corruptions. Second, we ablated ORA’s spatial masking and feature binding mechanisms and found the former to be most important for the accuracy of ORA’s predictions and the latter to be most important for the speed (i.e., number of iterations) that ORA took to make its recognition judgment. Third, we collected reaction time (RT) data from humans performing the same recognition tasks and found that, on trials having the longest RTs, ORA also needed more reconstruction-based feedback before recognizing the corrupted digit. In a fourth analysis we found that ORA and humans also shared the same pattern of errors (i.e., the same digit confusions). Lastly, we conducted a limited evaluation of ORA’s generalization to natural images by using the ImageNet-C dataset (Hendrycks & Dietterich, 2019), which consists of various corruptions but now applied to ImageNet objects. Details about the datasets and experimental procedures can be found in the Methods.

### 3.1 ORA improves recognition performance under visual corruptions

We used the MNIST-C dataset (Mu & Gilmer, 2019) to evaluate the effectiveness of ORA’s reconstruction-based feedback in improving recognition robustness under visual corruptions. This dataset includes 15 types of corruptions: affine transformation, noise, blur, occlusion, and others (See Extended Data S3 for example images of each). We also report results separately on a subset of the dataset referred to as **MNIST-C-shape**, which includes those corruptions relating mainly to object shape. Given ORA’s use of object-centric feedback, we hypothesized that robustness benefits would be greatest for these shape-specific corruptions. As shown in Table 1, this hypothesis was largely confirmed. We report results for two types of CNN model backbones used by ORA to approximate the features extracted by ventral visual brain areas, one a shallow 2-layer CNN (2 Conv) and the other a deeper 18-layer CNN with skip connections (Resnet-18; He et al., 2016). We also implemented three baseline models for comparative evaluation. Two were CNNs having the same deep (Resnet-18) or shallow (2 conv) backbones as ORA, but with added fully-connected layers for digit classification. For statistical analyses, we only compared the better performing of the two CNN baselines to ORA, which from Table 1 was the shallower 2 Conv CNN. We therefore refer to the better-performing 2 Conv baseline as “our CNN” for the remaining analyses. The third baseline was a CapsNet model (Sabour et al., 2017). Details about all three baselines can be found in the Methods.

**Table 1.** Model comparison results using MNIST-C and MNIST-C-shape datasets. Recognition accuracy (means and standard deviations from 5 trained models, hereafter referred to as model “runs”) from **ORA** and two **CNN** baselines, both of which were trained using identical CNN encoders (one a 2-layer CNN and the other a Resnet-18), and a **CapsNet** model following the implementation in Sabour et al. (2017).

| Dataset | ORA |  | CNN |  | CapsNet |
| --- | --- | --- | --- | --- | --- |
|  | Resnet-18 | 2 Conv | Resnet-18 | 2 Conv |  |
| MNIST-C | **91.84 (0.67)** | 88.32 (0.34) | 81.94 (0.43) | 88.94 (0.64) | 75.14 (1.00) |
| MNIST-C-shape | **96.24 (0.74)** | 92.82 (0.90) | 86.43 (0.44) | 91.80 (0.88) | 81.87 (0.55) |

The best recognition accuracies on both MNIST-C and MNIST-C-shape were from ORA with a Resnet-18 encoder. Moreover, during piloting we observed that using only the low-spatial frequencies in the image to generate the object reconstruction was sufficient to produce recognition robustness comparable to what is achieved with full-resolution object reconstruction (Extended Data S4). This shows that effective attention biases can be mediated by low-frequency neural feedback (Bar et al., 2006). ORA outperformed our CNN baseline by a significant margin (*t* (8) = 7.026, *p <* 0.001, *Cohen’s d* = 4.444), particularly excelling on the MNIST-C-shape dataset where it showed an improvement of over 4.5% (*t* (8) = 8.617, *p <* 0.001, *Cohen’s d* = 5.450). Despite incorporating object-centric representations for reconstruction (but lacking top-down attention feedback), the CapsNet model exhibited poor performance compared to ORA or our CNN baseline, indicating that the addition of reconstruction alone does not translate into improved accuracy and robustness. ORA was even more accurate than models specifically designed for generalization, such as CNNs trained using PGD/GAN adversarial noise (Extended Data S5). Lastly, we observed that increasing the capacity of the encoder from a 2-layer to an 18-layer CNN led to over-fitting by the CNN baseline and a consequent degradation of its accuracy from 88.94% to 81.94% on MNIST-C. ORA suffered no such degradation and achieved higher accuracy with the higher capacity encoder.

Figure 4A shows specific comparisons between ORA and our CNN baseline (the second best model) for each of the corruption types from MNIST-C-shape. For comprehensive results, refer to Extended Data S6. ORA consistently outperforms the CNN baseline and exhibits particularly strong performance under the fog corruption (*t* (8) = 5.318, *p <* 0.001, *Cohen’s d* = 3.363). As shown in Figure 4B, on average ORA confidently recognizes corrupted digits in 1.4 steps (measured as the number of forward processing steps required to reach a confidence threshold; see Methods). However, it takes significantly longer (more than 2 steps) under the fog corruption, suggesting that top-down attention feedback is particularly helpful in cases of visual corruptions that remove high-spatial frequencies.

**Fig 4.**
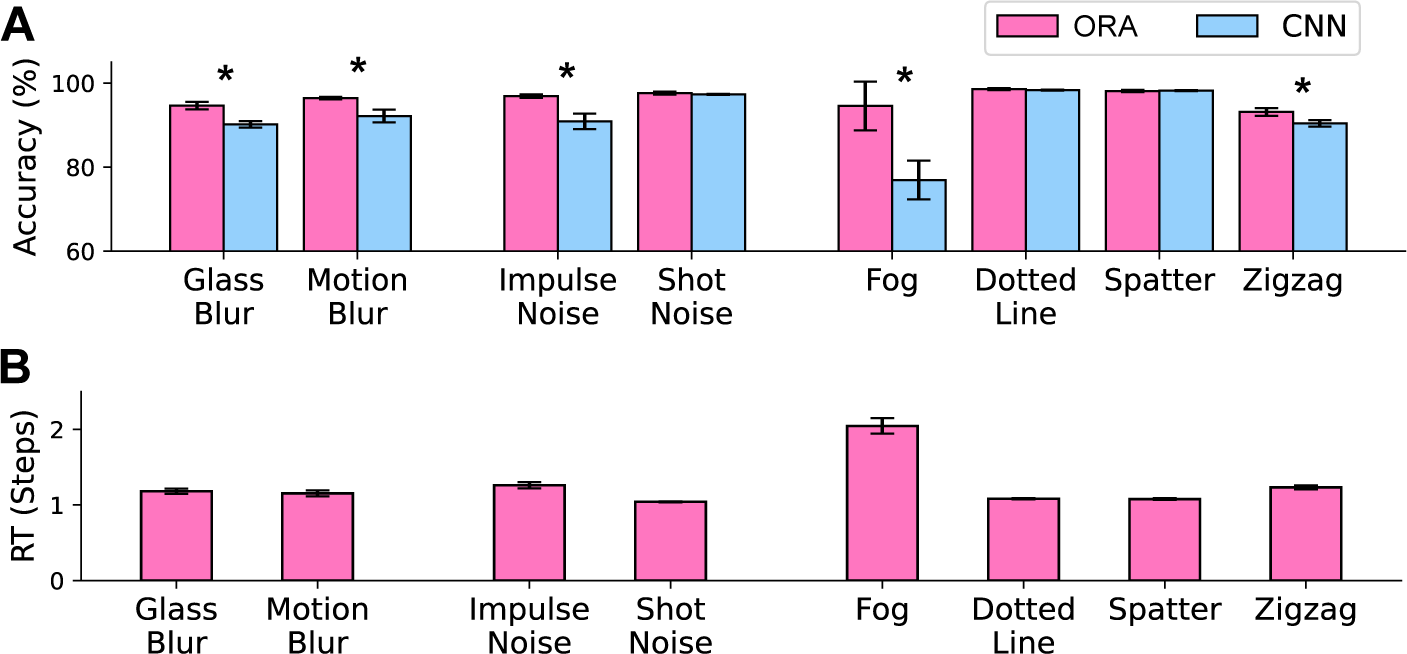
Model accuracy and reaction time for specific corruption types from MNIST-C-shape. **A:** ORA vs our CNN using each model’s best performing encoder (Resnet-18 and a 2-layer CNN, respectively). **B:** ORA’s reaction time (RT), estimated as the number of forward processing steps required to reach a confidence threshold. While many digits could be recognized in only one feed-forward pass, feedback mechanisms become useful for addressing specific types of noise, such as fog, or when the identity of the input is notably ambiguous, as demonstrated in Extended Data S8. Error bars indicate standard deviations from 5 different model runs. Asterisks (*) indicate statistical significance at a p-value of *<* 0.05.

### 3.2 Revealing distinct roles of spatial and feature-based attention

ORA uses two types of reconstruction-based feedback, roughly corresponding to spatial and feature-based attention, and we evaluated their unique contributions by ablating each from the best performing ORA model (i.e., Resnet-18 encoder) and observing the effect on performance. The left panel of Figure 5 shows that ORA’s recognition accuracy stemmed from its ability to use a generated reconstruction to perform spatial masking. Removing spatial masking led to a significant drop in performance of over 6% (-Spatial / +Feature; *t* (8) = 4.682, *p-adjust* = 0.009, *Hedges’ g* = 2.674). Ablating the feature binding component did not significantly affect prediction accuracy (+Spatial / -Feature; *t* (8) = 0.294, *p-adjust* = 1.0, *Hedges’ g* = 0.168), but this does not mean that it had no effect on performance. As shown in the right panel of Figure 5, feature-binding ablation instead lead to an increase in ORA’s RT (*t* (8) = 10.207, *p <* 0.001, *Cohen’s d* = 6.455). Note that we cannot report the corresponding spatial ablation because ORA’s RT is based on the number of spatial-mask iterations and would therefore always be zero. These findings suggest that ORA uses spatial and feature-based attention to perform distinct functional roles. Spatial attention improves recognition accuracy and enhances the neural signal-to-noise ratio by masking out noise from irrelevant spatial locations. On the other hand, feature-based attention enhances the overall discriminability of the encoder by suppressing features associated with competing object hypotheses. However, when the ventral encoder has a lower capacity for discriminability (i.e., the shallower 2-layer CNN), we observed that reconstruction-based feature binding also significantly influences both ORA’s RT and accuracy (Extended Data S7). The distinct roles played by feature and spatial attention in ORA’s function are further supported by the differential impact that each had on the model’s ability to recover from an initially incorrect hypothesis (Extended Data S1). On average, spatial attention led to an incorrect hypothesis changing to a correct prediction on approximately 6% of these trials, with a maximum of 40% incorrect-to-correct changes observed for the fog corruption. In contrast, iterations of feature-binding resulted in hypothesis changes on less than 1% of trials. This suggests that ORA uses feature-binding primarily to increase the efficiency of its recognition of MNIST-C digits whereas spatial masking is used to modify digit hypotheses and improve recognition accuracy.

**Fig 5.**
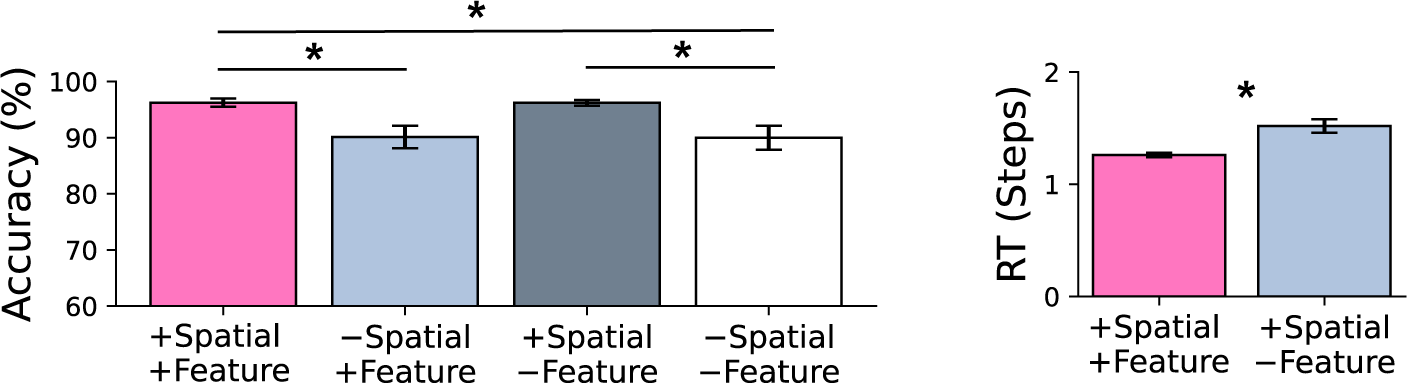
Results from ablating (-) ORA’s reconstruction-based spatial masking and feature binding components. Leftmost bars in pink indicate the complete (no ablation) ORA model. Left: Recognition accuracy. Right: reaction time, approximated by the number of forward processing steps taken to recognize digits with confidence. Error bars are standard deviations from 5 different model runs. Asterisks (*) indicate statistical significance at a p-value of *<* 0.05 with Bonferroni corrections.

### 3.3 Comparison with human reaction time

As shown in Figure 4B, ORA’s reaction time (RT) behavior changes depending on the difficulty of the recognition discrimination. The model iteratively generates the object reconstruction that it hypothesizes is the most plausible explanation of the input and uses it as a spatial mask for the subsequent feed-forward step. This process repeats until the model achieves a predetermined level of classification confidence measured by the entropy of the class likelihoods (softmax output). In the majority of cases, ORA reached the confidence threshold within a single feed-forward sweep (Figure 4B). However, on trials where the digit is inherently ambiguous or becomes ambiguous due to noise, ORA alternated between two object hypotheses, resulting in a longer time required for one to reach the confidence threshold (Extended Data S8).

To assess how this RT behavior of ORA compares to human RTs, we conducted a psychophysical experiment on humans who were asked to perform the same challenging MNIST-C recognition tasks (Figure 6A). Participants (N=146) viewed 480 images of digits and were instructed to press a keyboard spacebar as soon as they recognized the object in the image. Viewing time was response terminated and backward masked (see Methods for details). This RT judgment was followed by a second accuracy check in which participants used the keypad to enter the recognized digit. Only correct trials were used for the RT analysis. We selected MNIST-C digits for use as experimental stimuli based on the level of difficulty predicted by ORA, with our aim being to see whether human RTs will show a similar pattern. Specifically, we created easy, medium, and difficult conditions consisting of digits taking ORA either 1 step, 2-3 steps, or 4-5 steps to recognize, respectively (Figure 6B, left panel). Figure 6B (right panel) shows evidence for an alignment between human recognition accuracy and the three difficulty groups predicted by ORA. Human accuracy was by far the highest in the easy condition (i.e., only a single feed-forward step was needed by ORA for accurate model recognition), followed by the medium (2-3 steps) and difficult (4-5 steps) conditions, with the latter showing the most errors. The mean human RT was 1130ms (SD = 530ms), with a significant effect of condition difficulty, *F* (2, 284) = 105.266, *p <* 0.001, *η*^2^ = 0.426. Mean RTs in the difficult condition were 1265ms (SD = 649ms), but these decreased to 1170ms (SD = 517ms) in the medium condition and 985ms (SD = 473ms) in the easy condition. Lastly, Figure 6C shows a statistically significant positive Pearson correlation between human RTs and ORA’s RTs (*r* = 0.33, *p <* 0.001). This relationship extends our previous analysis of recognition difficulty by suggesting that ORA and humans found individual trials to be similarly difficult.

**Fig 6.**
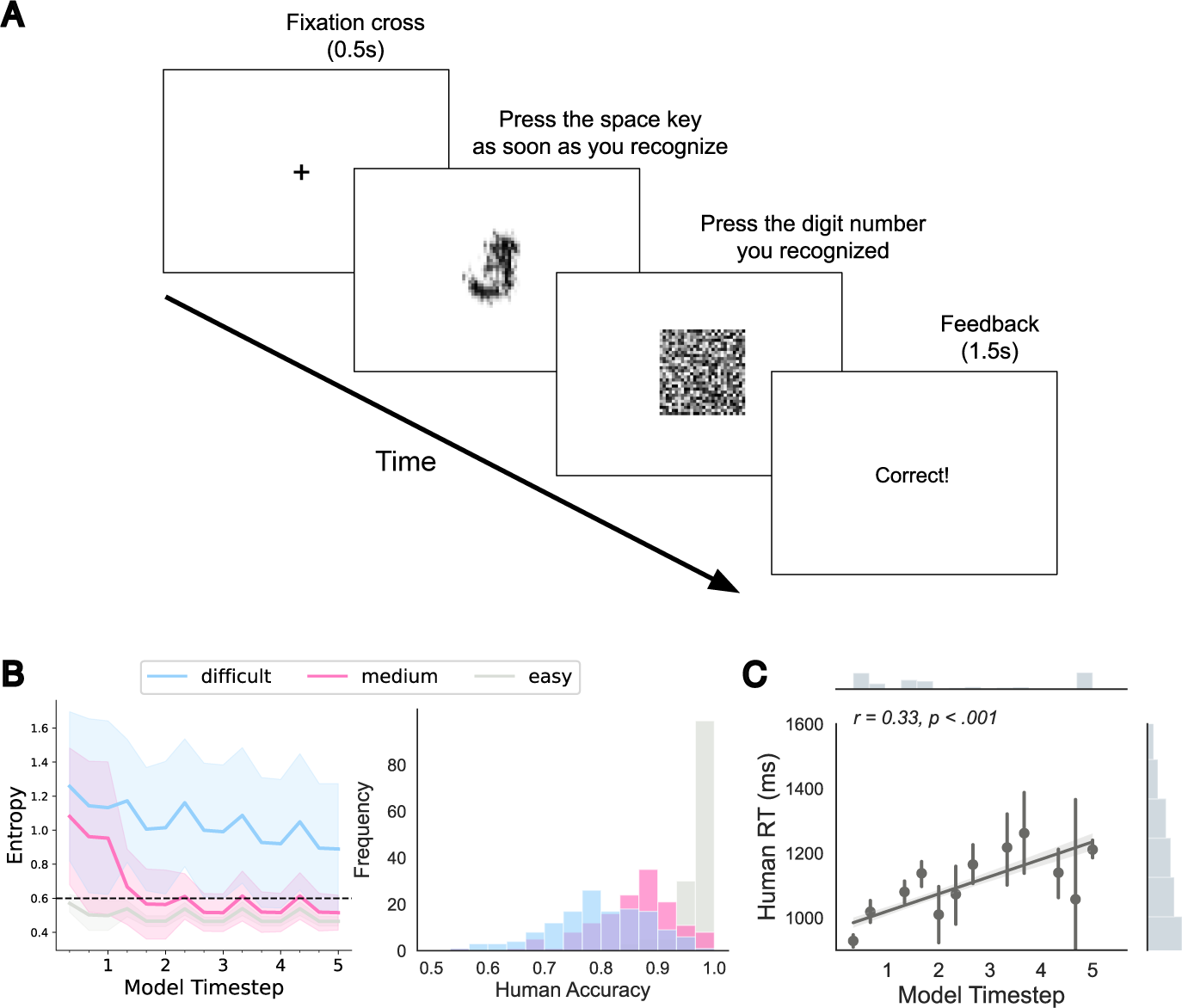
Human recognition experiment and results. **A**: Overview of the behavioral paradigm for measuring recognition reaction time (first button press) and accuracy (second button press). **B**: Stimuli were grouped into easy, medium, and difficult levels of recognition difficulty based on the number of forward processing steps needed for ORA to reach a predetermined recognition confidence threshold (dashed line; left panel). Colored regions indicate the corresponding standard deviations (SDs) across images. The unnumbered ticks on the x-axis indicate three local iterations of feature-based attention occurring within each forward processing (i.e., global spatial-masking iteration). The right panel shows distributions of human accuracy (averaged over participants) for the three difficulty conditions. **C**: Correlation between human RT and ORA’s RT, with the normalized marginal distribution of each variable shown on the top and right panels. Error bars represent standard error estimates bootstrapped from 1000 samples. Correlation plots for all MNIST-C corruption types can be found in Extended Data S9.

### 3.4 ORA’s errors and explain-away behavior

To gain a clearer insight into underlying processing mechanisms, here we make a deeper comparison between the errors generated by ORA and those from the best-performing CNN from Table 1. We found that ORA exhibits interesting systematic errors as it tries to explain-away random noise in the image and reconstruct it into a specific pattern that it has learned. For example, the model often connects noise speckles to generate a line, as demonstrated in the first row of Figure 7A, thereby mistakenly reconstructing a digit 9 instead of the ground truth digit 4. Similarly, ORA occasionally misinterprets zig-zag noise as being part of the digit, thereby reconstructing a digit 7 instead of the ground truth digit 1 (second row). These types of interpretable errors were not observed in the CNN baseline.

**Fig 7.**
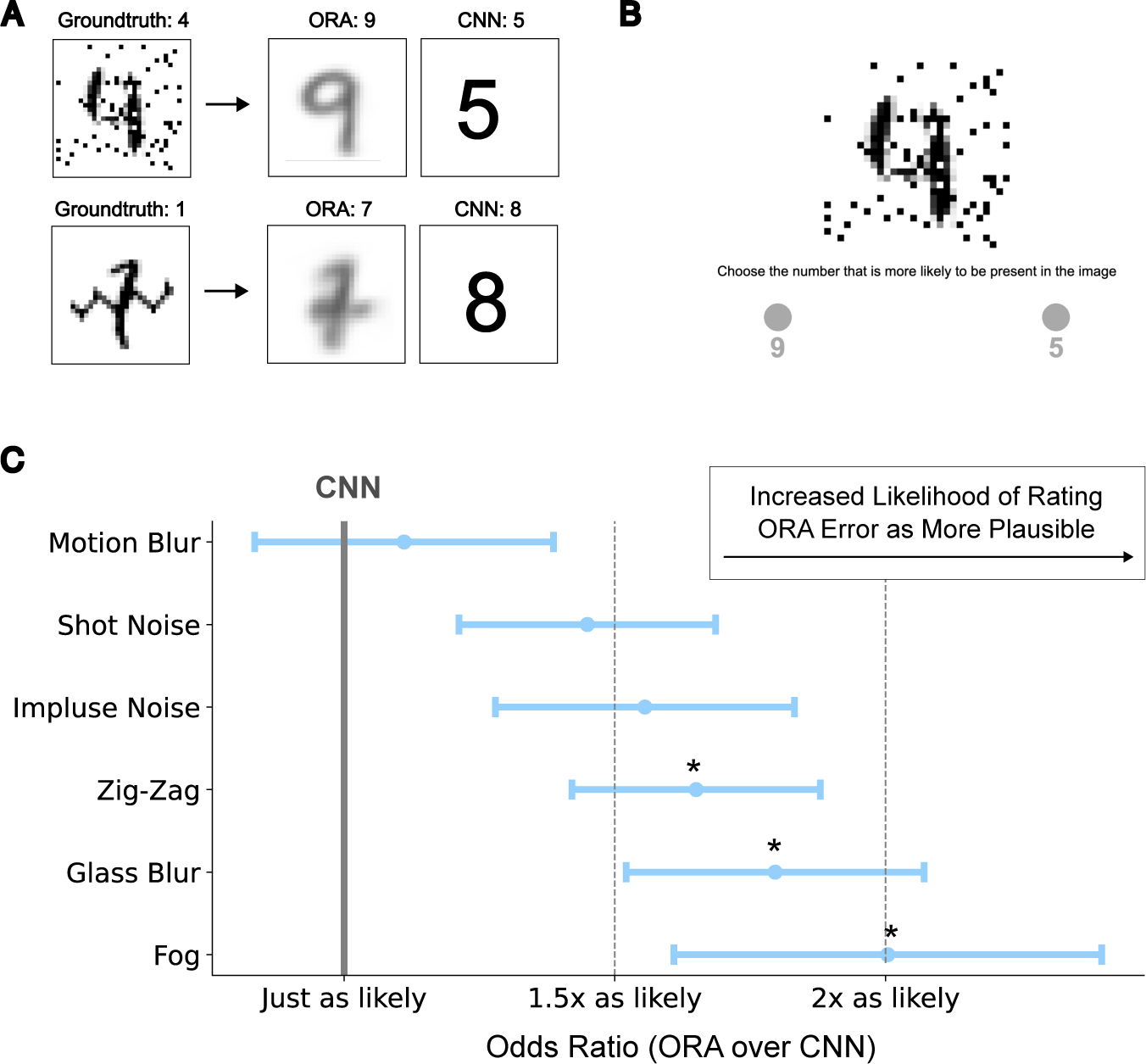
ORA often hallucinates a non-existing pattern out of noise, and these errors are perceived as more likely by humans compared to errors made by our CNN baseline. **A**: Example of ORA’s explain-way behavior and the resulting errors in classification. **B**: Two alternative forced choice experimental procedure requiring participants to choose which of two model predictions, one from ORA and the other from our CNN baseline (but both errors), is the more plausible human interpretation of the corrupted digit. **C**: The relative likelihood of ORA’s errors being perceived as more like those from a human, quantified as an odds ratio (a higher value indicates a more plausible ORA error). Error bars indicate standard errors, and asterisks (*) indicate odds ratios greater than 1 at a significance level of p *<* 0.05.

To determine whether human recognition errors are more aligned with ORA’s or our CNN baseline, we collected data from 184 participants in a two-alternative forced-choice task. On each trial, participants were shown a corrupted digit together with two mistaken model predictions, one from ORA and the other from the CNN, and they had to indicate which they thought would be more plausibly generated by a human (Figure 7B; see Methods for details). Note that we selected MNIST-C images as stimuli to satisfy two constraints: (1) that both ORA and our CNN baseline made an error in predicting the ground truth digit, and (2) that these two model errors differed from each other. To evaluate our data we computed an odds ratio, which measures the relative perceived likelihood of ORA’s errors compared to the CNN’s errors. We found that this ratio was consistently greater than 1 (with 1 indicating equally likely) for every type of corruption (Figure 7C). This was particularly true for the fog corruption, where ORA’s errors were rated twice as likely than those made by the CNN baseline (*t* (19) = 2.61, *p* = 0.02, *cohen’s d* = 0.58, testing against being *>* 1). Interestingly, the fog corruption type is also where ORA exhibited the highest performance advantage over the CNN (Figure 4). These results suggest that humans find ORA’s errors to be plausible, at least more so than errors from our CNN baseline.

### 3.5 Generalization to natural images

We evaluated whether ORA’s reconstruction-based feedback mechanism, shown to be beneficial for digit recognition, would generalize to more complex images of natural objects. We did this using the ImageNet-C dataset (Hendrycks & Dietterich, 2019), which is similar to MNIST-C but with visual corruptions applied to ImageNet objects (J. Deng et al., 2009). For model training we used a dataset called 16-class ImageNet (Geirhos, Temme, et al., 2018), which was developed to teach models a more human-like semantic structure by selecting from ImageNet a more comparable set of 16 basic-level categories (see Methods for the list of categories). To accommodate the complex backgrounds upon which objects appear in ImageNet images, we incorporated a recent segmentation model (Cheng, Schwing, & Kirillov, 2021) to estimate the ground-truth object boundaries, then trained the model to reconstruct only the pixels within the object segments. The resulting object reconstructions capture coarse representations of object shape, missing texture detail (see Extended Data S10).

Despite the coarseness of the generated reconstructions, their use as feed-back provided sufficient spatial masking to significantly improve recognition performance. Figure 8 reports model recognition accuracy (y-axes) for five increasing levels of corruption (x-axes) from the ImageNet-C dataset. We compared ORA’s performance to a CNN baseline model, now a ResNet-50 trained to recognize ImageNet objects. ORA’s feed-forward ventral process was approximated by the identical ResNet-50 (see Methods). Due to limitations in computational resources, we were able to conduct only a single training for each model (i.e., no error bars). Still, a clear pattern emerged. Although both models exhibit similarly accurate recognition at low corruption levels, at higher levels ORA is more robust on several types (on average *>* 7% than the CNN baseline without reconstruction feedback; Extended Data S11). The most notable benefits are observed with noise corruptions (top row), where accuracy increased for Gaussian noise by up to 35%. For some types of blur, the CNN baseline performed as well as ORA or slightly better, with the CNN working best for a pixelation corruption (bottom-right). These effects, however, are generally not as large as the improvements achieved by ORA (see Extended Data S11 for more results).

**Fig 8.**
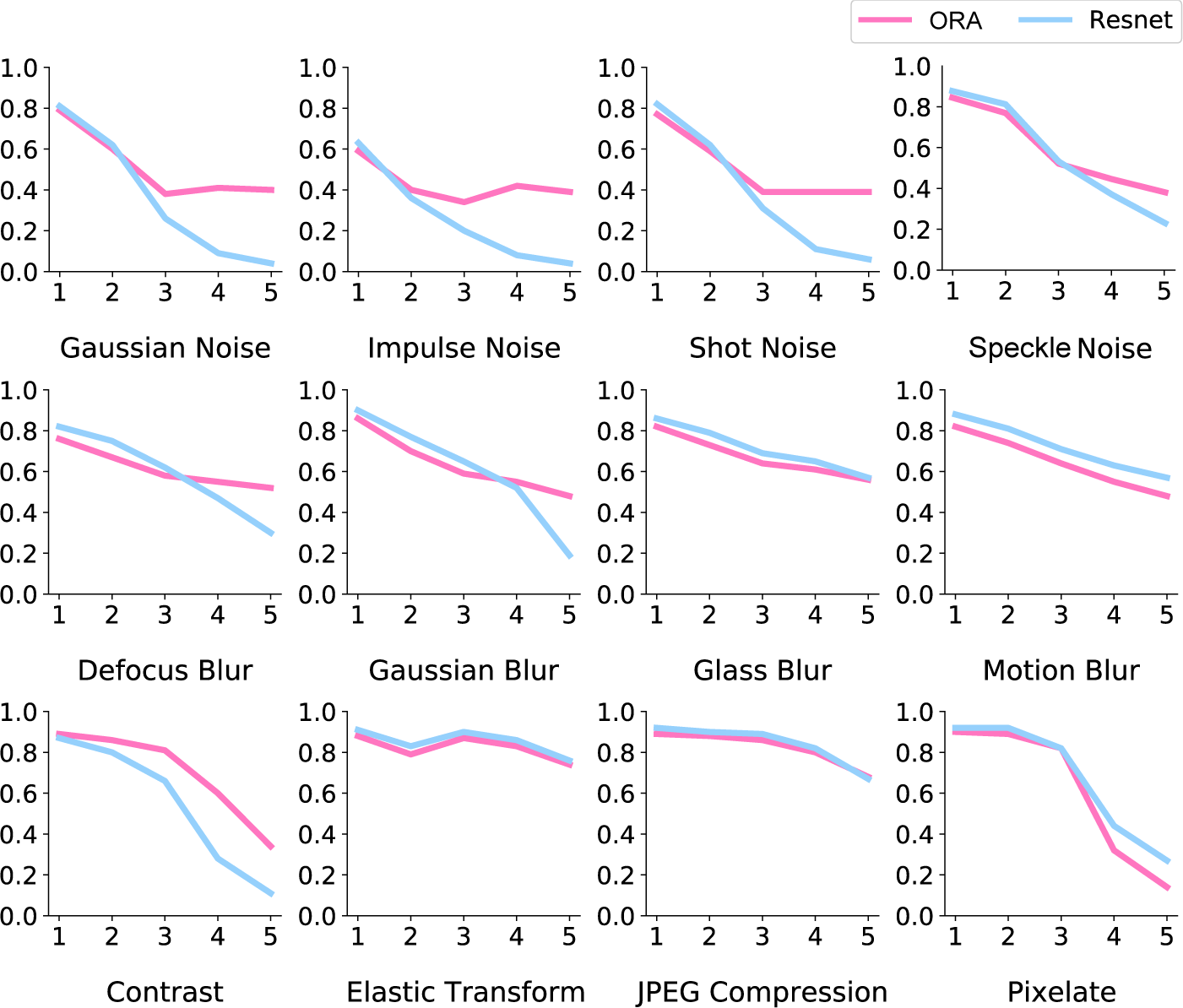
Model performance on the ImageNet-C dataset, with classification accuracy on the y-axes and level of corruption on the x-axes. ORA often demonstrates a clear advantage over the CNN baseline (ResNet-50), particularly with high levels of noise corruption.

## 4 Discussion

By whatever name is preferred, our common visual experience suggests that top-down, generative or reconstructive processes play a significant role in creating our visual perceptions. This is evident in phenomena such as visual imagery, object completion, and pareidolia. However, there are still open questions regarding *why* we have this ability to reconstruct objects and *what* functional role it plays in shaping our percepts (Breedlove, St-Yves, Olman, & Naselaris, 2020; DiCarlo et al., 2021; Gershman, 2019). Through a combination of computational modeling and behavioral experiments, we demonstrate that object reconstruction can serve as a mechanism for generating an effective top-down attentional template that can bias bottom-up processing to support challenging object recognition tasks. We implemented a neural network model of Object Reconstruction-guided Attention (ORA) that uses an autoencoder-like architecture (Hinton & Salakhutdinov, 2006; Sabour et al., 2017). ORA consists of an encoder that encodes the visual input into low-dimensional object-centric representations and a decoder that reconstructs objects from these representations. These object reconstructions are generated iteratively, enabling their use in routing the most relevant spatial and feature information to feed-forward object recognition processes. We found that ORA is highly robust in recognizing digits under a wide range of corruptions, outperforming other feed-forward model baselines. Theoretically, the functions performed by ORA’s reconstruction-based attention mechanism are similar to functions performed by spatial and feature-based attention (Fazl et al., 2009; Treisman, 1988). Empirically, we demonstrated that ORA is more predictive of human RTs and error patterns in recognition tasks compared to our CNN baselines. Lastly, we showed that the benefits of ORA’s reconstruction-based attention mechanisms generalize to recognition tasks using natural objects, with larger benefits observed in more challenging recognition tasks.

ORA shares many parallels with other influential cognitive neuroscience theories, including Mumford’s pattern theory (Mumford, 1994), Bar’s top-down prediction model (Bar et al., 2006), and other modeling approaches relying on interacting top-down and bottom-up processes (Carpenter & Grossberg, 1987; Dayan et al., 1995; Friston, 2010; Lee & Mumford, 2003; Ullman, 1995; Yuille & Kersten, 2006). Among these models, ORA builds most closely on adaptive resonance theory (ART; Carpenter & Grossberg, 1987). Both ORA and ART employ top-down templates to identify matches within bottom-up signals, selectively routing features that conform to these top-down expectations. However, a key difference lies in the formulation of the top-down templates. In ART, these templates are categorical and similar to prototypical representations of a given category (Minda & Smith, 2001). In contrast, ORA creates a top-down template from the visual input by using its generative attention mechanism to reconstruct an object category. It combines an autoencoder-like architecture with object-centric feature encapsulations to generate individualized top-down templates that are reconstructions from the individual image inputs, thus allowing for instance-specific features to be incorporated into categorical templates. This capability enables ORA to make detailed predictions about hypothesized objects and to facilitate precise attention control over low-level features—a critical missing link in the modeling of object-based attention (Cavanagh et al., 2023). ORA’s iterative and reconstruction-guided attention control over low-level features enables it to adjust its confidence in hypotheses, correct its errors, and explore alternative hypotheses without needing explicit “resets” or relying on memory-based inhibition, as implemented in ART.

One question not answered by our current work is how completely an object must be reconstructed in order to serve as an effective attentional template. Our finding that even low-resolution object reconstructions are sufficient to generate attentional biases and performance benefits align with previous studies showing that the brain initially processes low spatial frequency information about objects for the purpose of generating top-down feedback signals for guiding the bottom-up processing of finer-grained details (Bar, 2003; Bar et al., 2006; Bi, 2021). However, effects of feature-based attention are also found in a range of specific low-level features (e.g., color, orientation, etc, Maunsell & Treue, 2006), making further computational experiments needed to determine whether optimization of the information and level of object reconstruction might yield even greater performance benefits. Relatedly, although ORA’s current high compression rate limited its ability to reconstruct fine-grained object detail, it may be possible to address this limitation by exploiting recent advancements in decoder architecture and composition-based methods in the computer vision literature to generate reconstructions of varying kind and quality (Ho, Jain, & Abbeel, 2020; Singh, Deng, & Ahn, 2022). These questions, and broader questions regarding the biological plausibility of a reconstruction-based modeling approach, would benefit by additional study of the neural mechanisms underlying representation generation and object reconstruction in the brain. Studies on visual imagery have shown substantial overlap in neural processing between perception and mental imagery (Breedlove et al., 2020; Dijkstra, Bosch, & van Gerven, 2019), and more comparisons revealing similarities and differences between patterns of brain activity evoked by seen images and imagined images could provide valuable insights into the nature of the reconstruction process.

ORA has close theoretical ties to object-based attention mechanisms in that it uses object-centric representations (i.e., slots) to create top-down attentional biases (Cavanagh et al., 2023; Egly, Driver, & Rafal, 1994; Scholl, 2001; Vecera, 2000). However, while theories and models of object-based attention (Logan, 1996; Roelfsema & Houtkamp, 2011) have focused on the low-level grouping mechanisms responsible for combining visual elements into object parts and complete objects (e.g., gestalt rules; Wagemans et al., 2012), our generative approach highlights the role of top-down feedback in the creation of object entities. Specifically, we showed how top-down object-based feedback can mediate the selective binding of features and locations into objects, thereby implementing a form of object-based attention by integrating spatial and feature-based attentive processes (Kravitz & Behrmann, 2011; Scolari, Ester, & Serences, 2014; Shomstein & Behrmann, 2006). ORA accomplishes this by using top-down generative feedback to produce the most plausible object appearance given a visual input; it selects image locations and features that align most closely with the reconstructed object, a process biasing low-level neural responses to accurately reconstruct the input object. This fills a gap in the modeling of object-based attention by providing a mechanistic explanation of how high-level top-down expectations, such as object representations in the inferotemporal cortex (IT), can modulate low-level bottom-up visual features (Cavanagh et al., 2023).

Object-based attention has been studied using tasks ranging from object recognition to grouping to multiple-object tracking (Cohen & Tong, 2015; Roelfsema & Houtkamp, 2011; Scholl, 2001). Our study focused on object recognition under conditions of occlusion and noise, making our work more aligned with the use of object-based attention to create robust object entities (e.g., basic feature binding and grouping processes; Hummel & Biederman, 1992; Treisman, 1988) and less aligned with work emphasizing the operations performed on these entities and their limitations (e.g., multi-object tracking). Given the wide range of ways that researchers have conceptualized object-based attention, an important future direction will be to extend ORA and its evaluation to object-based attention tasks involving multiple objects. One potential area of investigation is perceptual grouping, which involves segregating an object of interest from other overlapping objects or a complex background. In our preliminary work on this question, shown in Extended Data S12, we found that ORA is showing promising results in segregating overlapping digits. This means that ORA has the potential to demonstrate classic effects of object-based attention, such as an ability to attend selectively to spatially overlapping faces or houses (Cohen & Tong, 2015; O’Craven, Downing, & Kanwisher, 1999). Another interesting direction will be to apply ORA to the creation and maintenance of multiple objects in a scene. This task will require the model to maintain feature bindings for different objects belonging to either the same or different object categories, and is an essential step in extending ORA to multiple-object tracking. Such explorations would likely lead to dynamic slot representation and potentially reveal higher-level relationships between attention control and its modulation by short-term memory (Bundesen et al., 2011).

Our study has broad implications for the fields of behavioral vision science and computer vision. While there is still only limited empirical evidence for the use of generative processes in vision (Dijkstra & Fleming, 2023; Gershman, 2019), there is growing interest in capturing the robustness of human perception by modeling it as a reconstructive process (Adeli et al., 2023; Al-Tahan & Mohsenzadeh, 2021; Boutin et al., 2020; Csikor et al., 2022; Fleming & Storrs, 2019; Hedayati et al., 2022; Xing et al., 2022; Yildirim et al., 2020). A reconstruction framework offers the potential to unify our understanding of top-down cognitive processes, including attention biases (Cavanagh et al., 2023), memory retrieval (Hedayati et al., 2022), hypothesis testing (Bar et al., 2006), and mental imagery (Breedlove et al., 2020). We also hope that our work will inspire the development of novel computational architectures in computer vision. Methods that train models to reconstruct objects have been shown to enhance generalization performance (Dittadi et al., 2022; Locatello et al., 2020), but this approach has primarily been explored in the context of pretraining or incorporating auxiliary loss (He et al., 2022; Sabour et al., 2017). Our work is the first to show that a reconstruction approach can effectively model attentional biases during inference, which had been an open research question (Shi, Darrell, & Wang, 2023).

In summary, our study demonstrates that the attentive reconstruction of objects can serve as a plausible neural mechanism for achieving robust object perception and recognition. We model this mechanism as ORA, which uses object reconstruction as a top-down attention bias to focus bottom-up processing on information relevant to what it considers to be the most plausible (i.e., recognizable) object shape in the visual input. This object reconstruction-based mechanism unifies space-based and feature-based modes of attention into one computational attention system.

## 5 Methods

### 5.1 ORA Implementation

For ORA’s encoder, which models the ventral stream, we used different pytorch implementations of feedforward CNNs: one a shallow 2-layer CNN^2^ (2 Conv) and a deeper CNN^3^ with skip connections (Resnet with 18 and 50 layers for the MNIST-C and ImageNet-C tasks, respectively; He et al., 2016). For ORA’s decoder we used three fully-connected dense layers to reconstruct the MNIST image and an inversely mirrored Resnet to reconstruct the ImageNet images. As in the original CapsNet model (Sabour et al., 2017), we used a vector implementation of object and feature slots, where the length of each slot vector represents the presence of an object class in the image. To make this value range from 0 to 1 (i.e., 0 for absence and 1 for presence), we used the following nonlinear squash function. We also provide specification details for object slot representations (Sabour et al., 2017) and the total number of trainable parameters.

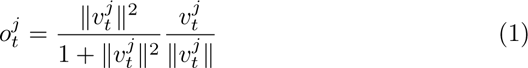

### 5.2 Loss Function

ORA outputs both object slot activations and image reconstructions, and the losses for each output are computed and combined using a weighting parameter of *λ_recon_* = 0.392 (Eq.2). For classification, we used margin loss (Sabour et al., 2017) to match the object slot activations to the ground truth (Eq. 3). The first term of the loss is active only when the target object is present (*T_j_ >* 0), and minimizing this term pushes the object slot activation (measured by the vector magnitude *||o_j_||*) to be closer to 1, with a small margin (*m*). The second term applies only when the target object is absent, and minimizing this term pushes the predicted object slot activation below the small margin (*m*).

|  | for MNIST-C task |  | for ImageNet-C task |
| --- | --- | --- | --- |
| Encoder type | 2 Conv | Resnet-18 | Resnet-50 |
| Num of feature slots | 1152 | 288 | 3136 |
| Feature slot dimension | 8 | 8 | 8 |
| Num of object slots | 10 | 10 | 16 |
| Object slot dimension | 16 | 16 | 49 |
| Total number of parameters | 2,904,720 | 4,581,296 | 67,116,160 |

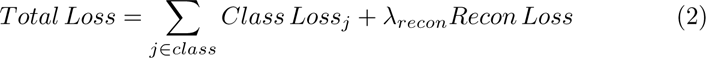

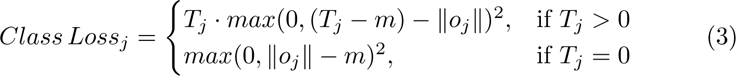

### 5.3 Baseline Implementation

The CNN baselines consist of the same encoders used for ORA (See Section 5.1) followed by two fully connected layers for classification readout. The readout layers are trained using the dropout technique for regularization (Srivastava, Hinton, Krizhevsky, Sutskever, & Salakhutdinov, 2014). Negative Log Likelihood loss is computed on the log softmax output for classification. The total number of model parameters are: 1,199,882 for the 2-layer CNN baseline, 2,803,882 for the ResNet-18 baseline in the MNIST-C experiment, and 23,540,816 for the ResNet-50 baseline in the ImageNet experiment. Performance for the other baselines reported in Extended Data are taken from the original benchmarking paper (Mu & Gilmer, 2019).

### 5.4 Training Details

Both ORA and our CNN baselines were trained and validated on clean images (MNIST, L. Deng, 2012 for a digit recognition task; ImageNet, J. Deng et al., 2009 for a natural object recognition task) and tested using a separate dataset containing unseen images subjected to various types of corruptions (MNIST-C, Mu & Gilmer, 2019 for a digit recognition task; ImageNet-C, Hendrycks & Dietterich, 2019 for a natural object recognition task). These separate training and testing datasets ensure that models are evaluated on their ability to generalize their learned object representations to previously unseen exemplars and types of corruption, a challenge commonly known as the out-of-distribution problem in computer vision (Geirhos et al., 2021). All model parameters were updated using Adam optimization (Kingma & Ba, 2014) with a mini-batch size of 128 for the MNIST-C experiment and 32 for the ImageNet-C experiment. For the MNIST-C experiment, we used the exponentially decaying learning rate by a factor of 0.96 at the end of each training epoch, starting from 0.1. Similarly, for the ImageNet-C experiment, the learning rate followed the same exponential decay pattern, but with a starting value of 0.0001. Model training was terminated based on the results from the validation dataset (10% of randomly selected training dataset) and was stopped early if there was no improvement in the validation accuracy after 20 epochs. Models were implemented in the PyTorch framework and trained on a single GPU using a NVIDIA GeForce RTX 3090 (24 GB) for the MNIST-C experiment and a RTX A6000 (48 GB) for the ImageNet-C experiment.

### 5.5 Measuring ORA’s Reaction Time

ORA’s processing is limited to five forward processing steps, each followed by reconstruction-based spatial attention feedback (i.e., global spatial masking iteration). However, the model typically did not need all five of these forward processing steps in order to confidently recognize an object, and the number of steps actually taken by the model to reach a classification threshold is ORA’s RT. This threshold is measured by entropy over the softmax distribution of class likelihoods, similar to the method used in Spoerer, Kietzmann, Mehrer, Charest, and Kriegeskorte (2020), and we chose an entropy threshold of 0.6 based on a grid search. If this threshold is not reached within the five forward processing steps, we use the prediction from the final time step for classification.

### 5.6 Spatial Masking Methods

In each forward processing step, the model generates a reconstruction-guided spatial mask based on the most likely object reconstruction. To create this mask, we simply apply a threshold to the reconstructed pixel values. We used thresholds of 0.1 for MNIST images and 0.05 for ImageNet images, where the pixel intensities in both range from 0 to 1. This thresholding process creates a boolean mask where pixels with values below the threshold are assigned 0 and pixels with values above the threshold are assigned 1. The masked input for the next forward processing step is obtained by element-wise multiplication of the reconstruction with the boolean mask. The non-zero areas of the masked input are normalized using MaxMin normalization (Equation 4, when *p* corresponds to the reconstructed pixel value at image coordinates *i* and *j*). For ImageNet images, the RGB reconstructed images are first converted to grayscale before applying the thresholding and normalization processes.

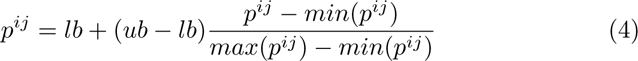

### 5.7 Feature Binding Algorithm

ORA’s reconstruction-guided feature binding process computes coefficients to dynamically modulate weights between features and objects. These coefficients, denoted as *C_ij_*, are computed iteratively over three local iterations occurring within each forward processing step (i.e., global spatial masking iteration). We update the original weights by multiplying them by the final coefficients, *C_ij_ ·W_ij_*. In the original CapsNet model (Sabour et al., 2017), these binding coefficients are modulated based on representational similarity, which is calculated as the dot product between the features and objects (Line 6 in the Pseudocode). In our model, we also adjust these binding coefficients to suppress the features that are connected to objects having high reconstruction error. By suppressing the features leading to inaccurate object reconstructions, those features that are most selective to the hypothesized object become bound. All coefficients are normalized over objects using MaxMin normalization (like Equation 4).

**Algorithm 1.**
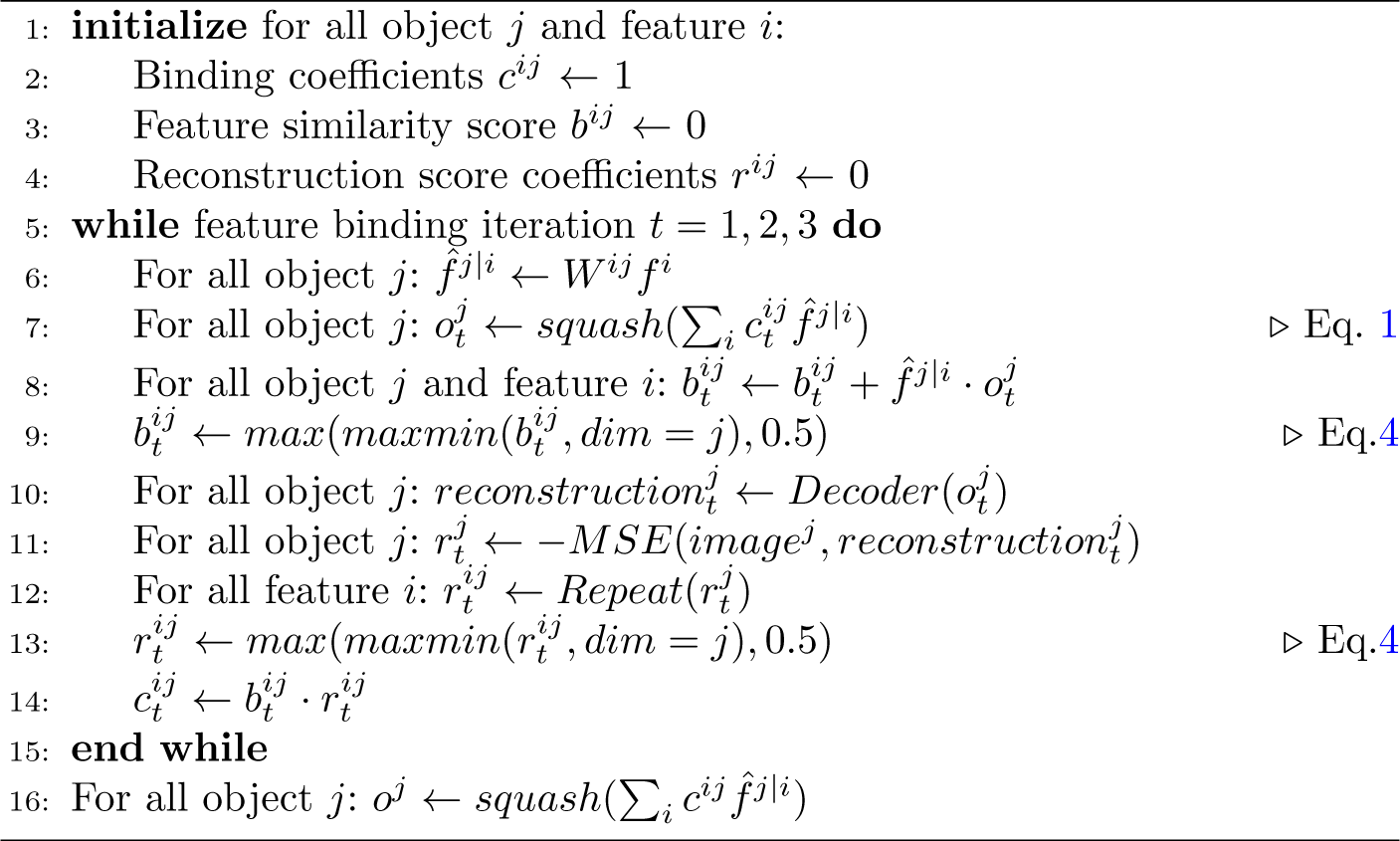
Reconstruction-guided Feature Binding.

### 5.8 16-Class-ImageNet Training Dataset

The original Imagenet dataset includes 1,000 subordinate-level labels. These include 120 different dog breeds, which results in a semantic structure that is highly atypical compared to that of an average person (Ahn, Zelinsky, & Lupyan, 2021) that may influence model performance. To address this problem, we also trained ORA using 16-class-ImageNet, a subset of ImageNet that groups its 1000 fine-grained classes into 16 basic-level categories: airplane, bicycle, boat, car, chair, dog, keyboard, oven, bear, bird, bottle, cat, clock, elephant, knife, and truck (Geirhos et al., 2021; Geirhos, Temme, et al., 2018).

### 5.9 Behavioral Methods for the MNIST-C Recognition Experiment

#### Participants

A total of 146 Stony Brook University undergraduates, self-identified as 94 women, 47 men, and 2 non-binary individuals, participated in the experiment for course credit. This sample size was determined based on an a priori power analysis using G*Power (Faul, Erdfelder, Lang, & Buchner, 2007), which indicated that a minimum of 28 participants would be sufficient for a repeated-measures ANOVA to detect a medium effect size with a power of .80 at a significance level of *α* = .05. This estimate is based on each participant viewing a 96 image subset of the 480 total testing images (see Stimuli and Apparatus), a design choice motivated by a desire to avoid participant fatigue and to maintain their engagement in the task. Participants all had normal or corrected to normal vision, by self report. This study was approved by the Stony Brook University Institutional Review Board, and written consent was obtained from all participants.

#### Stimuli and Apparatus

Stimuli consisted of 480 images from the MNIST-C testing dataset Mu and Gilmer (2019), 60 images for each of the 8 corruption types in MNIST-C-shape. Images were divided into 5 sets of 96 images, where each participant viewed a 96 image set. To experimentally test ORA’s RTs we selected the images used for behavioral testing based on the RT predicted by the model. Specifically, for each corruption type we selected 20 images by randomly sampling from 3 different model RT conditions: short (1 step), medium (2-3 step), and long (4-5 step). Note that all images were correctly classified by the model, meaning that this manipulation is targeted to the number of forward processing steps needed by ORA to recognize an object. Selected images were presented on a 19-inch flat-screen CRT ViewSonic SVGA monitor at a screen resolution of 1024*×*768 pixels and a refresh rate of 100 Hz. Participants were seated approximately 70 cm from the monitor and at this viewing distance the image height subtended approximately 8*^◦^*of a visual angle. Space bar responses were made with the participant’s dominant hand using a standard computer keyboard.

#### Procedure

Participants were instructed that they would see a series of centrally presented digits, ranging from 0 to 9, and that they should press the space bar as soon as they recognized the digit class. However, the task emphasized accuracy as well as speed. Participants were instructed to make their key press when they felt confident in their answer, although a reminder was provided if the response was too slow (*>* 5 sec). Upon pressing the space bar, the original stimuli was masked by a randomly generated pixel image and participants were asked as part of a secondary judgment to report the digit that they recognized by pressing the digit on the numeric keypad. At the start of each trial a central cross appeared immediately prior to stimuli presentation (500 ms), and feedback was provided at the end of each trial on the accuracy of the digit recognition response (10 alternative forced choice). Each participant viewed 96 images, 4 images *×* 3 model RT conditions *×* 8 corruption types. Ten practice trials were provided before the experiment so that participants could familiarize themselves with the task. The order of presentation of the images was randomized over participants.

### 5.10 Behavioral Methods for the MNIST-C Error Likelihood Rating Experiment

#### Participants

A total of 184 undergraduate students from Stony Brook University, self-identified as 120 women, 61 men, and 3 non-binary individuals, participated in the experiment for course credit. Sample size was determined based on an a priori power analysis using G*Power (Faul et al., 2007), which indicated that a minimum of 41 participants per stimulus set (4 stimulus sets in total) would be sufficient for a paired t-test to detect a medium effect size with a power of .80 at a *α* = .05 level of significance. All participants had normal or corrected to normal vision, by self report. This study was approved by the Stony Brook University Institutional Review Board, and written consent was obtained from all participants.

#### Stimuli and Apparatus

Stimuli consisted of 120 images selected from the MNIST-C testing dataset Mu and Gilmer (2019), 20 images per corruption type, to satisfy the experimental constraint of having competing models make different errors to a stimulus. These ambiguous images were divided into 4 separate sets, with each set containing 30 images. Only six corruption types from the eight MNIST-C-shape categories were used because the number of images meeting the experimental constraint (i.e., different errors) were too few for analysis in the dotted-line and spatter corruption conditions. The stimuli image spanned 40% of their screen size in height. All responses were made with a computer mouse or trackpad.

#### Procedure

Participants accessed the rating experiment remotely using their desktop devices. On each trial, a single corrupted digit was presented at a size of 40% of the vertical screen dimension. Below this stimulus were the two model predictions, horizontally separated (the location was randomized), and participants were instructed to rate which of the two would be more plausibly made by a person (two alternative forced choice; see Fig. 7B). A response was required on each trial and participants were not allowed to skip images, although they could change their answers anytime before proceeding to the next trial (which they initiated by clicking a “Next” button). Three practice trials were provided before the experiment so that participants would familiarize themselves with the task and interface. Each participant viewed 30 images. The order of presentation of the images was randomized over participants.

## Code availability

The code necessary to reproduce our results is available on the github repository at https://github.com/ahnchive/ORA-recognition

## Supporting information

**S1 Table.**
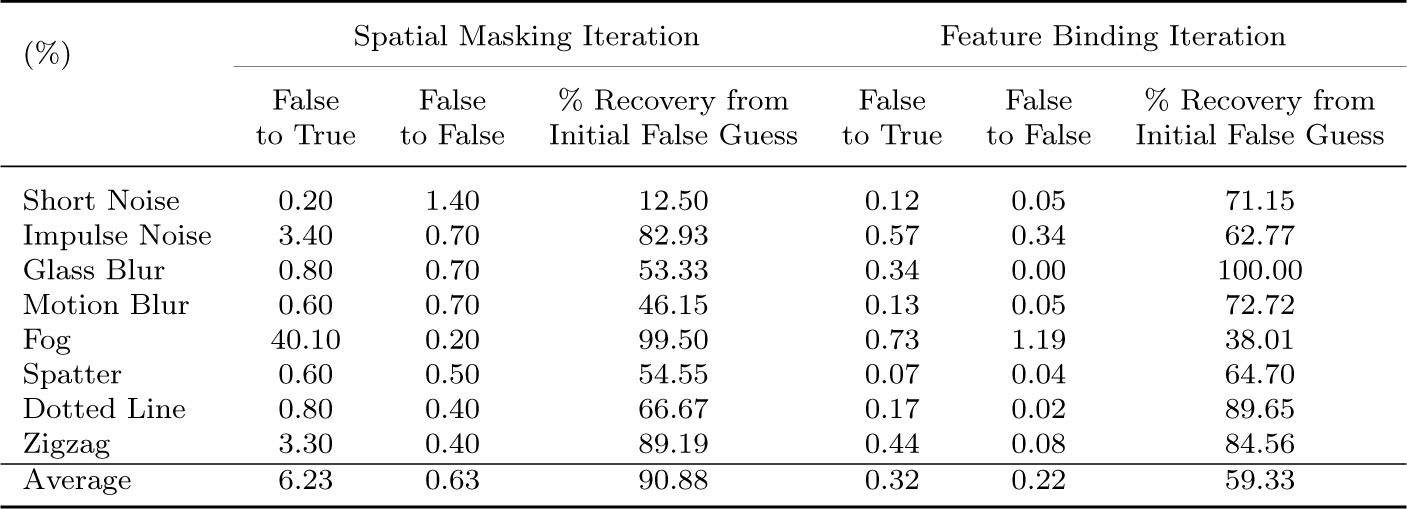
Percentage of changes in ORA’s class prediction during spatial masking and feature binding iterations.

S1 Percentage of changes in ORA’s class prediction during spatial masking and feature binding iterations
| (%) | Spatial Masking Iteration |  |  | Feature Binding Iteration |  |  |
| --- | --- | --- | --- | --- | --- | --- |
|  | False to True | False to False | % Recovery from Initial False Guess | False to True | False to False | % Recovery from Initial False Guess |
| Short Noise | 0.20 | 1.40 | 12.50 | 0.12 | 0.05 | 71.15 |
| Impulse Noise | 3.40 | 0.70 | 82.93 | 0.57 | 0.34 | 62.77 |
| Glass Blur | 0.80 | 0.70 | 53.33 | 0.34 | 0.00 | 100.00 |
| Motion Blur | 0.60 | 0.70 | 46.15 | 0.13 | 0.05 | 72.72 |
| Fog | 40.10 | 0.20 | 99.50 | 0.73 | 1.19 | 38.01 |
| Spatter | 0.60 | 0.50 | 54.55 | 0.07 | 0.04 | 64.70 |
| Dotted Line | 0.80 | 0.40 | 66.67 | 0.17 | 0.02 | 89.65 |
| Zigzag | 3.30 | 0.40 | 89.19 | 0.44 | 0.08 | 84.56 |
| Average | 6.23 | 0.63 | 90.88 | 0.32 | 0.22 | 59.33 |

**S2 Fig.**
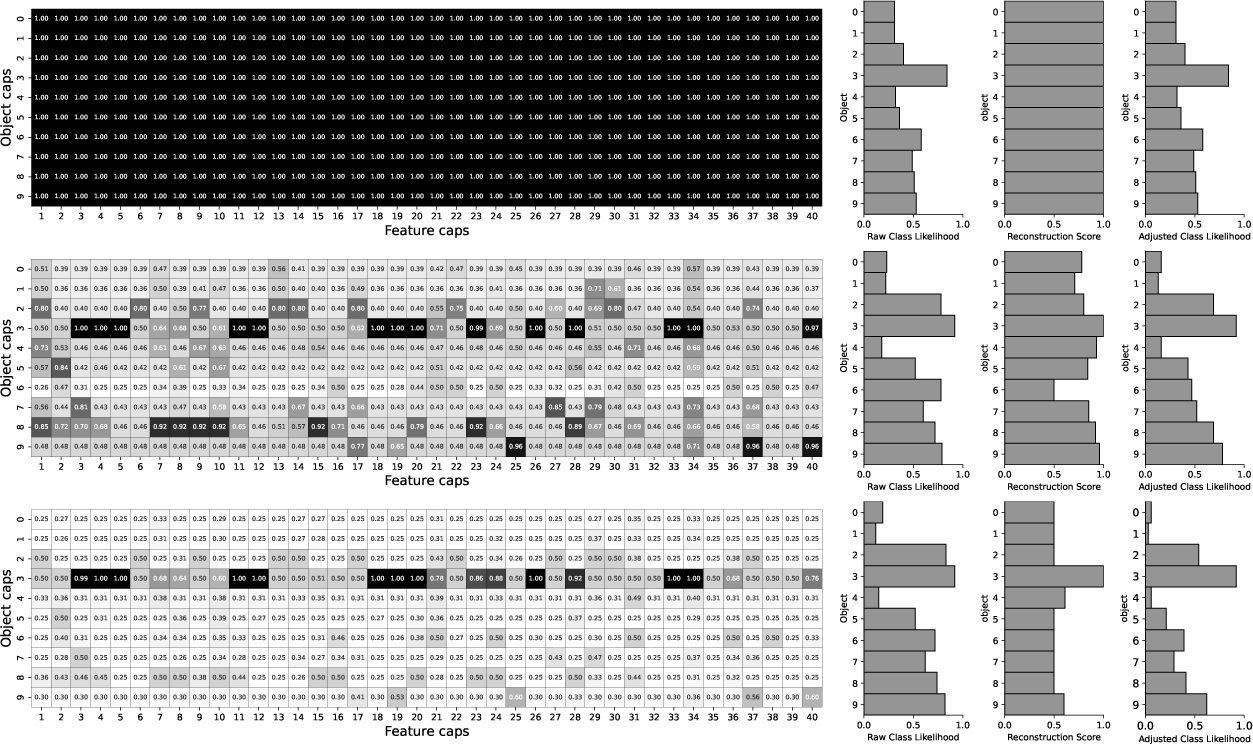
Step-wise visualization of reconstruction-guided feature binding process. The left column of the figure below shows binding coefficient matrices between all 10 of the possible digit categories (rows) and the first 40 feature slots (columns) at each of the 3 feature binding iterations. Binding coefficients are initially unweighted (step 1, top) but those object features leading to greater reconstruction error become suppressed (become lighter) over iterations, resulting in the feature binding coefficients focusing on the object features that best match the input (middle and bottom matrices). The nine bar plots in the right of the figure illustrate the effect of reconstruction-guided feature binding on ORA’s digit classifications. The leftmost three plots show the raw likelihood of classification for each of the 10 object slots at the start of each time step (i.e., before the application of reconstruction-guided feature binding). The middle three plots show reconstruction scores (inverse of reconstruction error) at each step, and the rightmost three plots show the updated class likelihood after reconstruction-guided feature binding is applied (i.e., the two are multiplied). Best viewed with zoom in PDF.

**S3 Fig.**
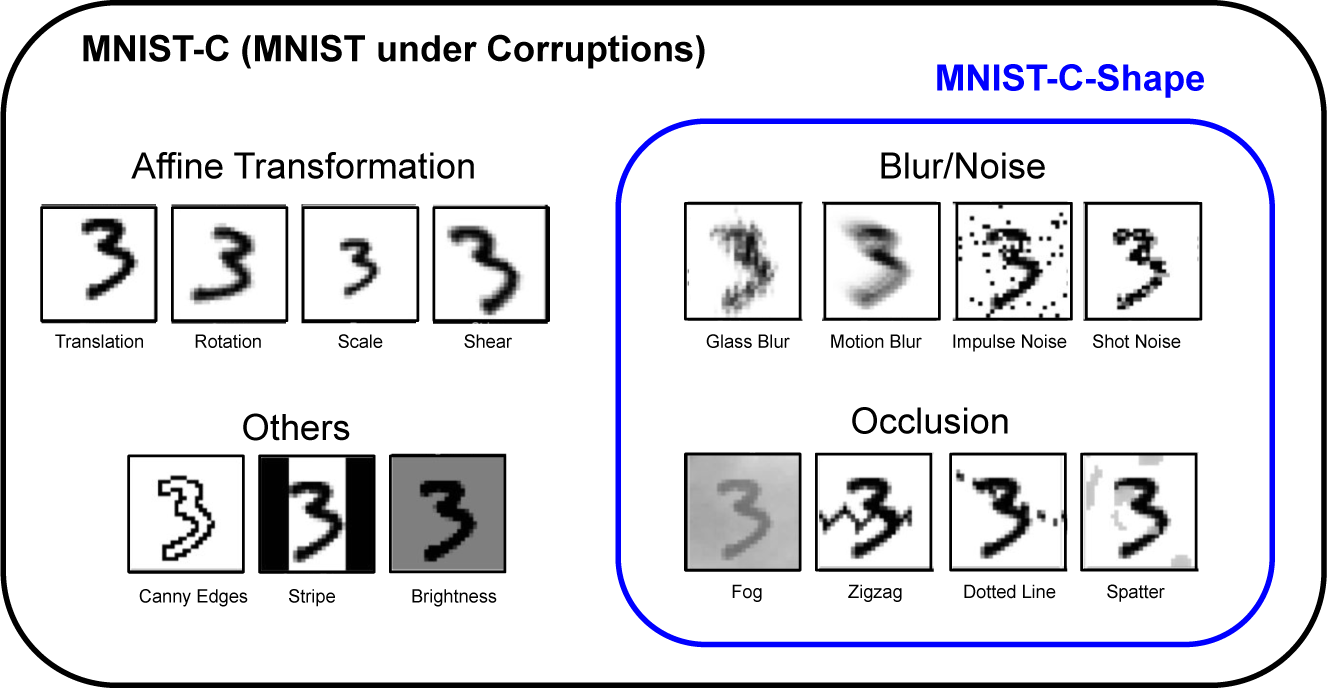
MNIST-C dataset examples.

**S4 Table.**
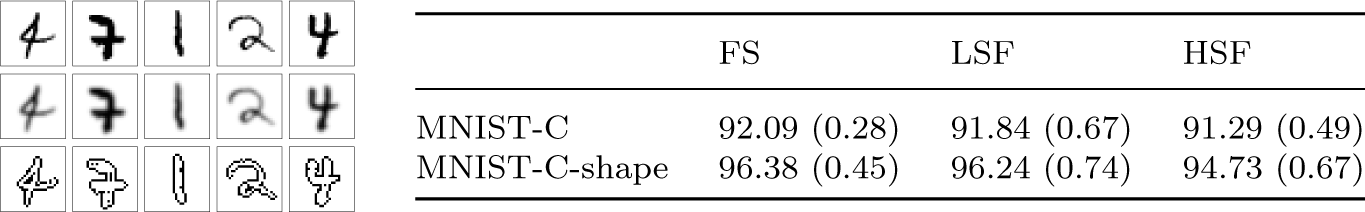
Results from manipulating the spatial frequencies used for image reconstruction. To test how reconstruction efficacy is effected by spatial frequency, we manipulated the spatial frequencies of the visual input that ORA used to generate its reconstructions. we trained different models to reconstruct different spatial frequency components of the input (e.g., generating a low spatial frequency reconstruction of the original full spectrum image) and then tested them to evaluate the effect of spatial frequency on model performance. We extracted different spatial frequency information from the input using a broadband Gaussian filter with cutoff frequencies below 6 cycles per image and above 30 cycles per image for generating low and high spatial frequency images, respectively. As shown below, we found that the use of a low-resolution object reconstruction yields comparable performance to a model trained with the full spectrum. The figure on the left shows samples of reconstructions from full-spectrum images (top) and images filtered for low (middle) and high (bottom) spatial frequencies. The table on the right shows the average performance of models trained to reconstruct either full-spectrum (FS), low-spatial frequency (LSF), or high-spatial frequency (HSF) images. Values in parentheses indicate standard deviations from 5 model runs.

**S5 Table.**
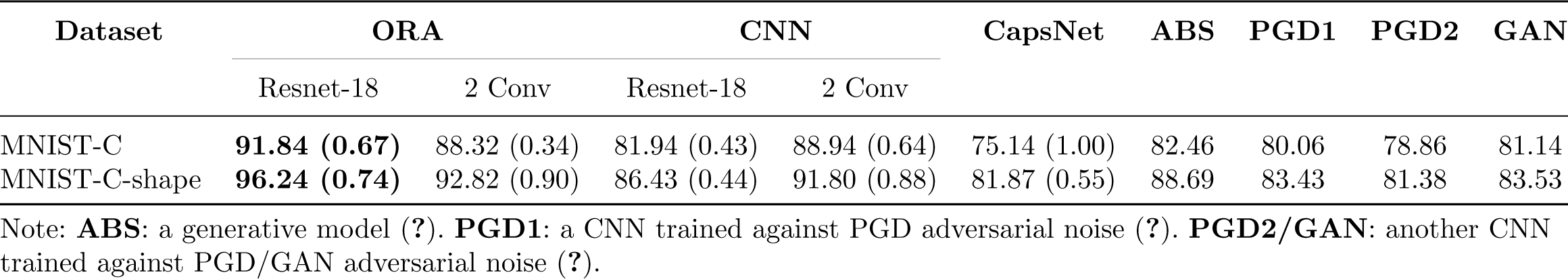
Model performance comparison. Extended model comparison. Average performance and standard deviations from 5 model runs are reported for **ORA**, **CNN**, and **CapsNet** models that we trained for this study (see text for details). Best model performance for the other baselines are taken from Mu and Gilmer (2019).

**S6 Fig.**
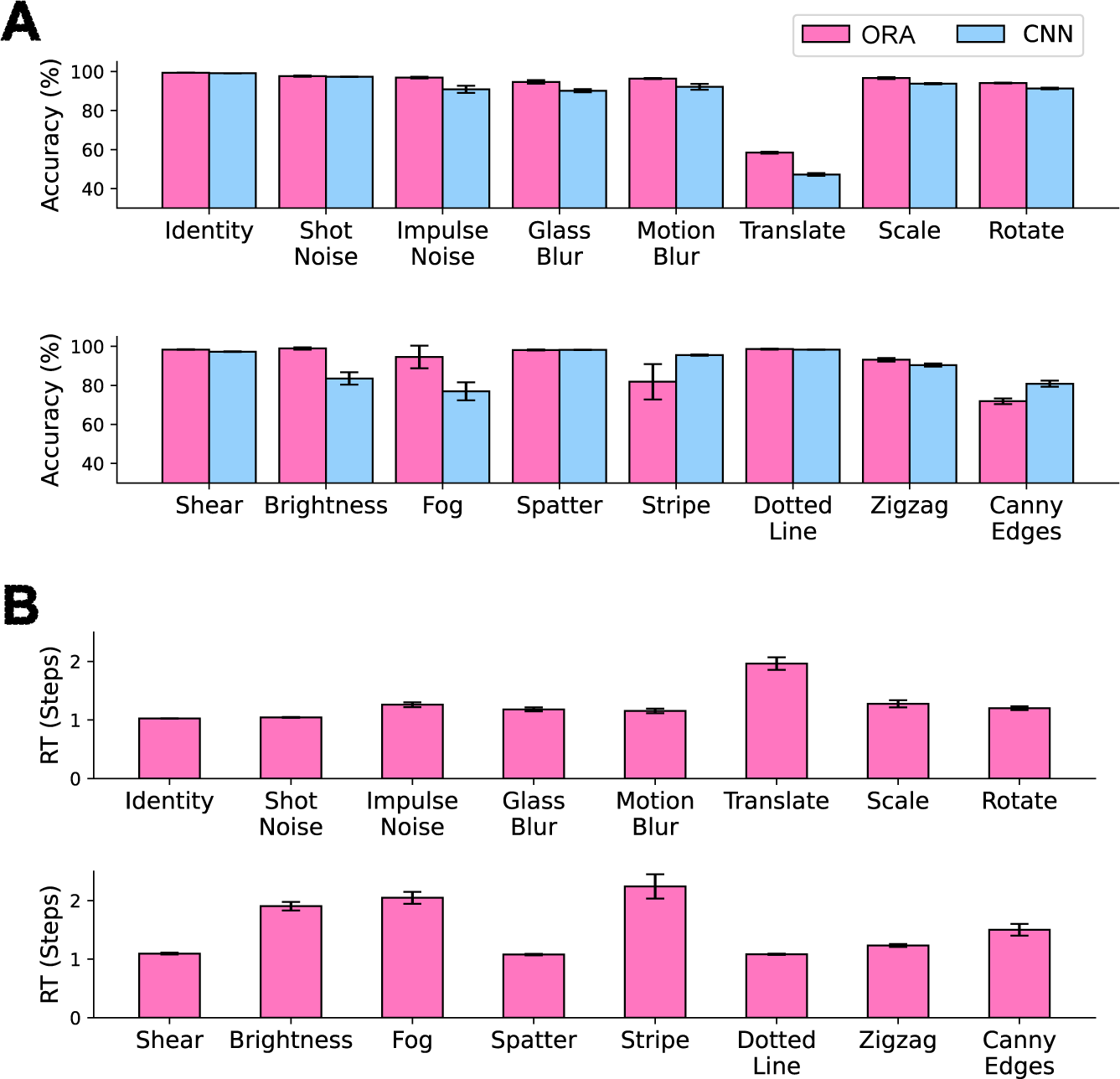
Model comparison on all MNIST-C corruption types. Average model accuracy and reaction time for all of the MNIST-C corruption types. A: ORA and CNN accuracy using each model’s best performing encoder (Resnet-18 and a 2-layer CNN, respectively). B: ORA’s reaction time (RT), estimated as the number of forward processing steps required to reach a confidence threshold. Error bars indicate the standard deviations from 5 different runs.

**S7 Fig.**
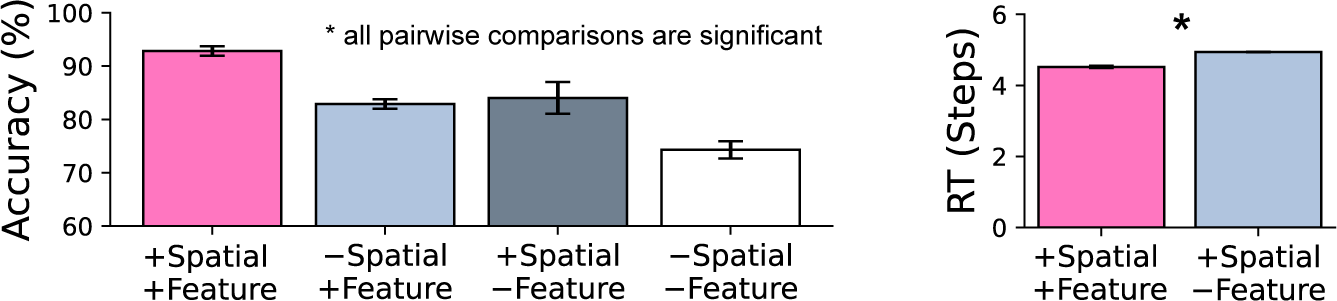
Ablation studies on ORA with 2 Convolutional encoder. Results from ablating ORA’s reconstruction-based spatial masking and feature binding components. Ablated components are indicated by a - symbol, none ablated components are indicated by a +. Leftmost bars in pink indicate the complete (no ablation) ORA model. Left: Recognition accuracy. Right: reaction time, approximated by the number of forward processing steps taken to recognize digits with confidence. Error bars are standard deviations from 5 different runs.

**S8 Fig.**
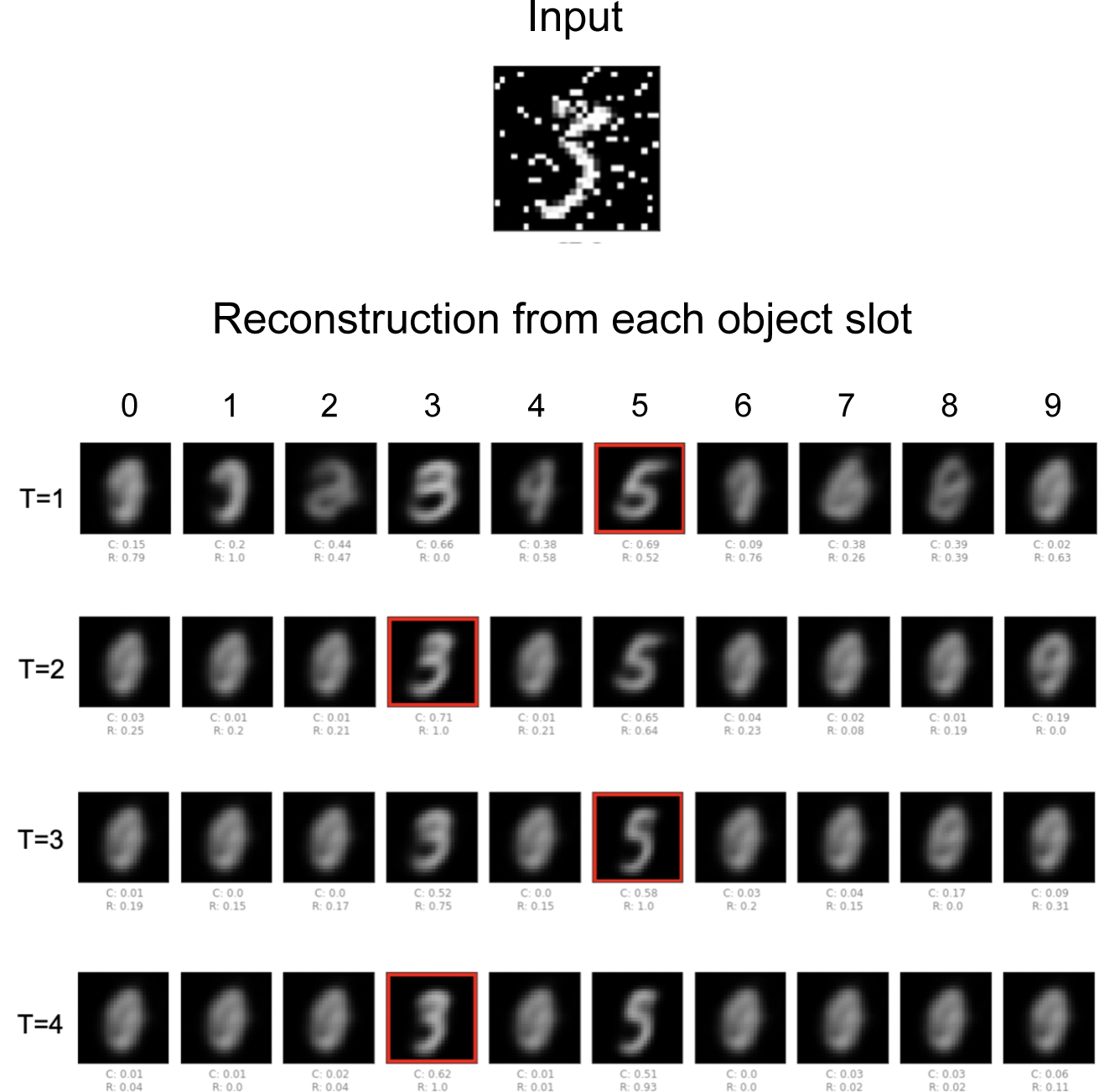
ORA’s behavior on an ambiguous stimulus. The figure below shows ORA taking longer to converge on a single digit hypothesis when the input is ambiguous. On the top is the input image, which could be plausibly interpreted as either the digit 3 or the digit 5. On the bottom are reconstructions hypothesized for all 10 digits, with the respective classification (C) and reconstruction (R) scores below each. ORA’s most likely object hypothesis (with the highest class likelihood) at each step is indicated by the red square. Note that ORA’s best guess about this object vacillates between the 3 and 5 hypotheses over time steps. The ground truth for this image is the digit 3.

**S9 Fig.**
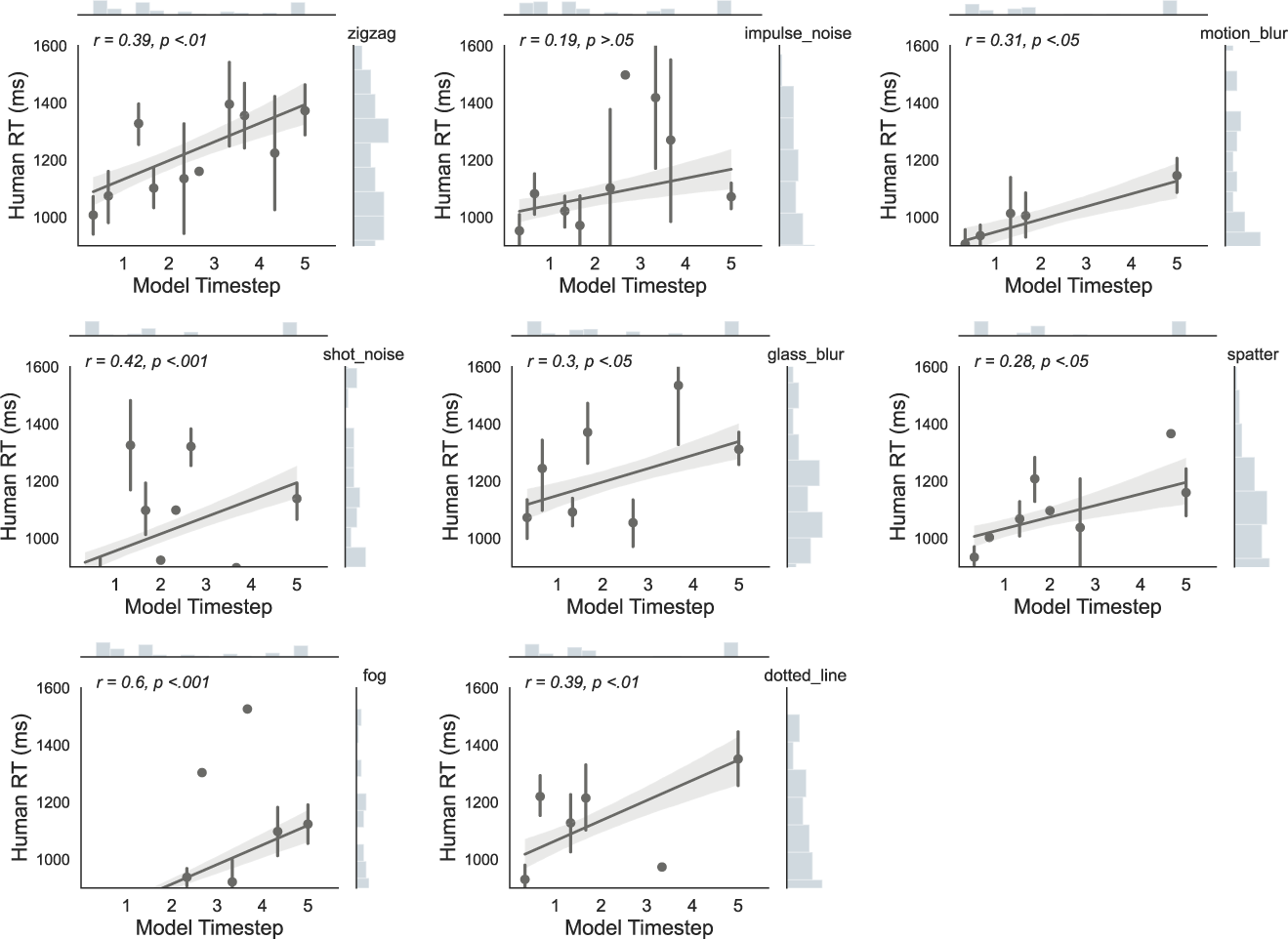
Correlations between human and ORA reaction times for all MNIST-C corruption types.

**S10 Fig.**
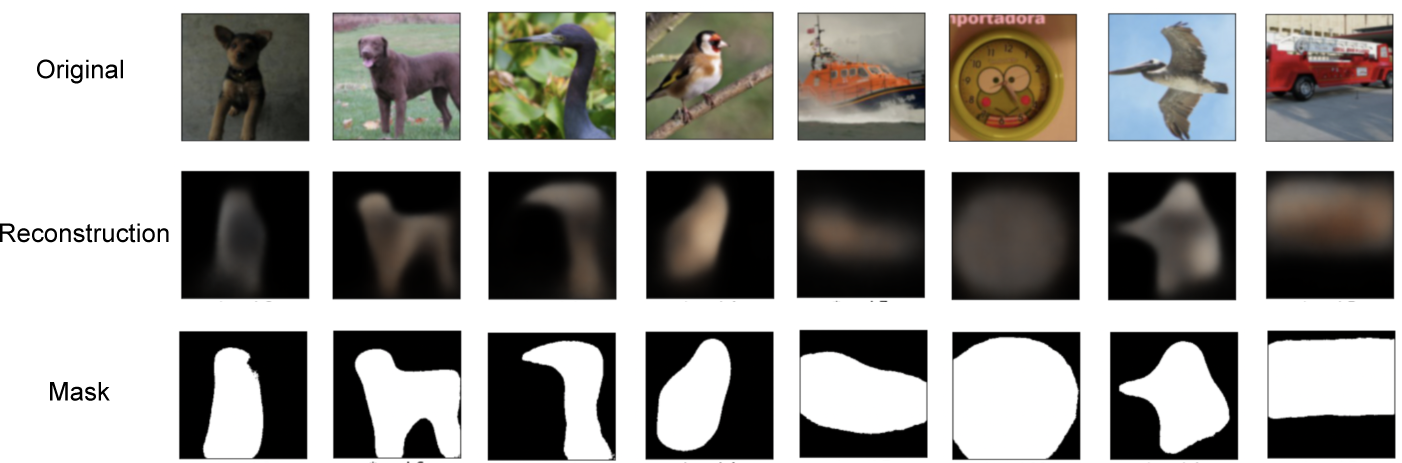
Examples of ORA reconstructions and spatial masks from Imagenet-C.

**S11 Fig.**
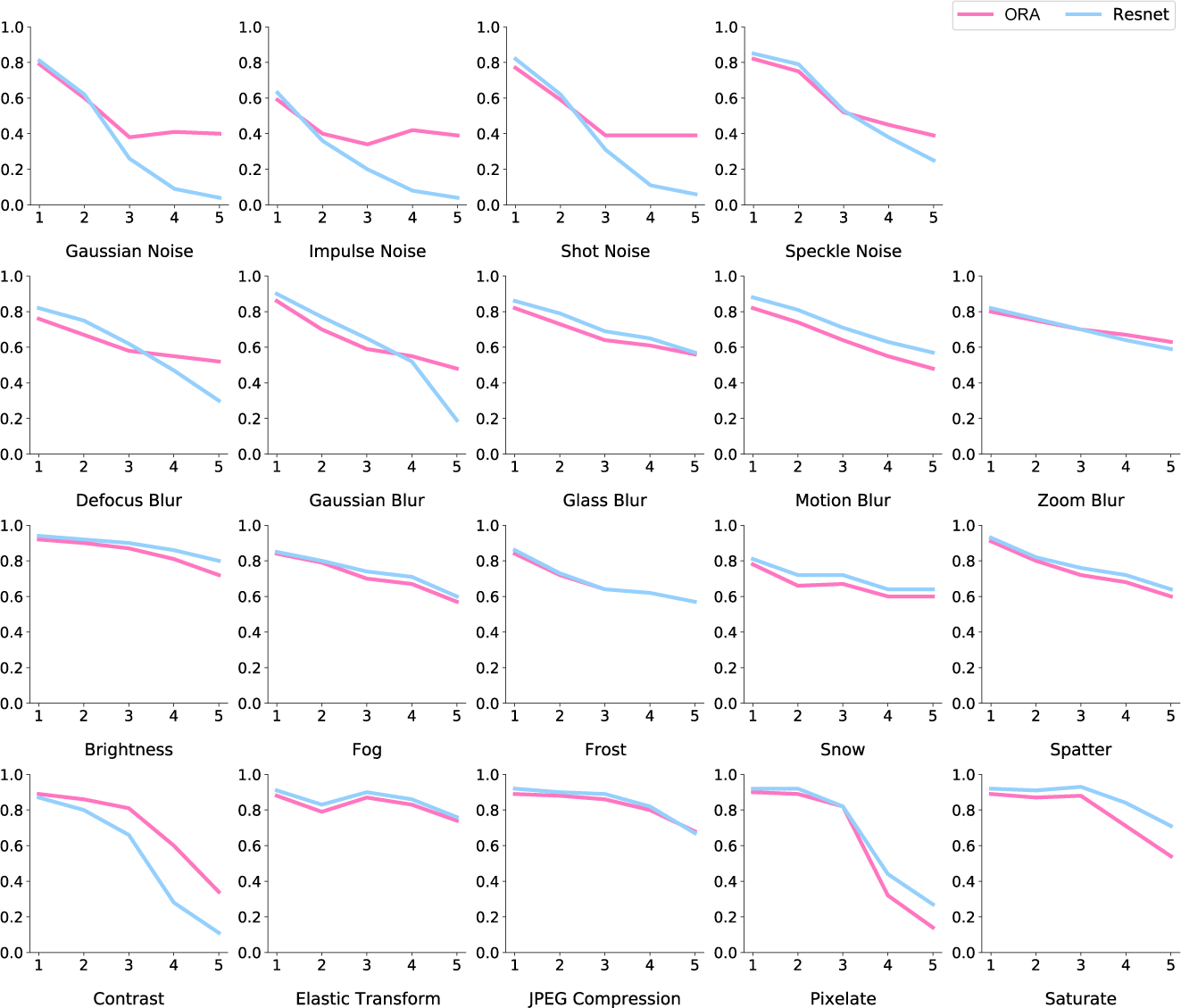
Model comparison for all ImageNet-C corruption types. The x-axes show five level of noise severity and the y-axes show classification accuracy.

**S12 Fig.**
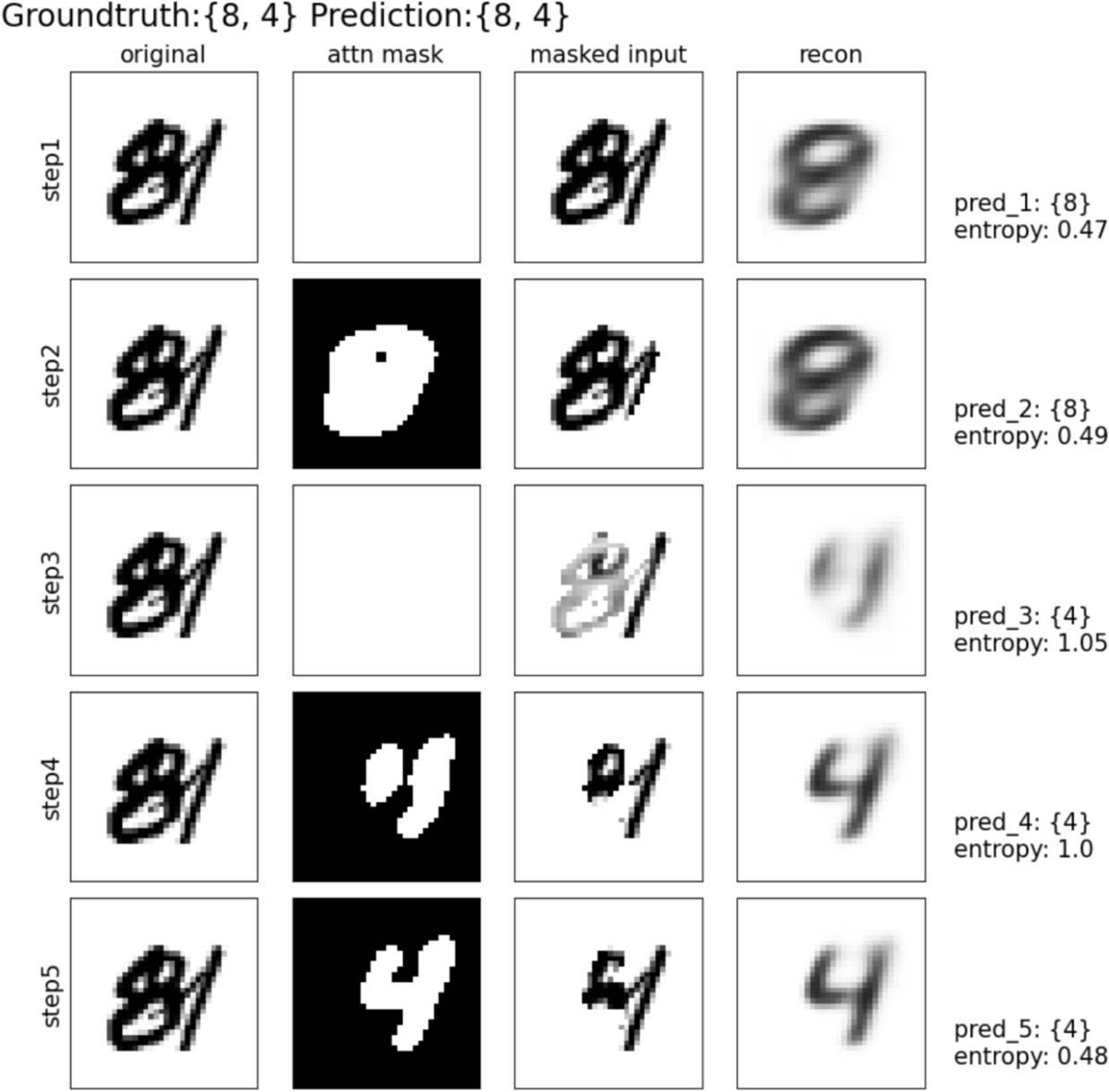
Applying ORA to an overlapping-object recognition task. To apply ORA to an overlapping-digit recognition task we generated the MultiMNIST dataset as described in Sabour et al. (2017), where each image consists of two MNIST digits overlapped, randomly shifted 1-4 pixels horizontally and vertically in opposite directions. The resulting images are 36 × 36 pixels, with an average 70% overlap between digit bounding boxes. A total of 10,000 images were randomly selected from the MNIST testing dataset for model evaluation. The figure shows a representative overlapping-digit stimulus and model outputs illustrating how ORA can solve the overlapping-digit task. The ORA model utilized in this demonstration is identical to the one presented in our paper, meaning it was trained for single-digit recognition and reconstruction, and we performed no additional hyperparameter tuning. The only change that we made in our application of ORA to overlapping-digit recognition was algorithmic; once the model recognizes a first digit with sufficient certainty (using an entropy threshold of 1.0 as a measure of confidence), it proceeds to identify the next digit. It does this by expanding its spatial mask and focusing on the input signals that remain unexplained by first digit recognition. This approach therefore allows the model to sequentially attend to individual overlapping objects, effectively segregating and recognizing each one after the other, which parallels the seriality observed in human recognition (Broadbent, 1957; Pashler, 1998). In the illustrated example, the ground truth consists of the digits 8 and 4 overlapped. ORA initially identifies the digit 8 before shifting its attention to 4, repeating this process until the confidence threshold is achieved. Using its sequential object attention mechanism, ORA outperforms the best-performing CNN baseline, a ResNet-18 (a shallower 2-layer CNN performed poorly on this task). Prediction accuracy was measured using two metrics: image-level accuracy, where the response is considered correct only if both objects in an image are identified accurately, and object-level accuracy, which reflects the average number of objects recognized (0.5 indicates one of two objects are recognized, while 1.0 indicates both objects are recognized). ORA’s predictions are generated from the first two digits recognized with high confidence within a maximum of 10 forward iterations. The CNN baseline generated predictions using the top 2 most activated class outputs. Although there is room for improvement, this initial application of ORA to overlapping digits is already beating a strong baseline recognition model.

| (%) | Our<br>w/o Feedback | Our<br>w/Feedback<br>(Spatial Only) | Our<br>w/Feedback<br>(Spatial & Feature) | CNN baseline |
| --- | --- | --- | --- | --- |
| Image-level Accuracy | n/a | 36.54 | **41.01** | 18.10 |
| Object-level Accuracy | 39.97 | 61.34 | **64.32** | 51.33 |

## Footnotes

1 In computer vision, this encapsulation of information is sometimes referred to as a “capsule” (Doerig, Schmittwilken, Sayim, Manassi, & Herzog, 2020; Sabour, Frosst, & Hinton, 2017), although the term that we adopt is the more recently used “slot” (Greff, van Steenkiste, & Schmidhuber, 2020; Locatello et al., 2020). Note that in cognitive psychology, a slot corresponds to one “file” of information about an object as theorized by object file theory (Kahneman, Treisman, & Gibbs, 1992; see also the idea of category-consistent features in Yu, Maxfield, & Zelinsky, 2016).

2 https://github.com/pytorch/examples/blob/main/mnist/main.py

3 https://github.com/pytorch/vision/blob/main/torchvision/models/resnet.py

